# The ribosome quality control factors Asc1 and Hel2 regulate the expression of HSP70 during heat shock and recovery

**DOI:** 10.1101/2022.09.12.507689

**Authors:** Lokha R. Alagar Boopathy, Emma Beadle, Alan Xiao, Aitana Garcia-Bueno Rico, Celia Alecki, Irene Garcia de-Andres, Maria Vera

## Abstract

Cells rapidly adapt to survive harsh environmental conditions through the potent upregulation of molecular chaperones or heat shock proteins (HSPs). The inducible members of the HSP70 family are the fastest and most transcriptionally induced chaperone upon stress. The *HSP70* mRNA life cycle regulation in the cytoplasm is unique because it is translated during stress when general translation is repressed and rapidly degraded once conditions are optimal for growth. Contrary to the role of the *HSP70* mRNA 5’ untranslated region in maximizing the synthesis of HSP70, we discovered that the coding sequence (CDS) represses its translation through the ribosome quality control (RQC) mechanism. The CDS of the most inducible HSP70 in *Saccharomyces cerevisiae, SSA4*, is uniquely biased with low-frequency codons that promote ribosome stalling during heat stress. The stalled ribosomes are recognized by RQC components Asc1p and Hel2p and two ribosome proteins, Rps28A and Rps19B, that we identified as new RQC components. Surprisingly, RQC does not signal the degradation of the *SSA4* mRNA by no-go-decay (NGD). Instead, Asc1p destabilizes the *SSA4* mRNA during recovery from heat stress by a mechanism independent of its ribosome binding and *SSA4* CDS codon optimality. Therefore, Asc1p operates two synergistic mechanisms that converge to regulate the life cycle of *HSP70* mRNA during stress and recovery. Our research identifies Asc1p as a critical regulator of the stress response and RQC as the system tuning HSP70 synthesis.

## INTRODUCTION

Cells have mechanisms to mitigate the detrimental effects caused by stressors in the surrounding environment, such as an increase in temperature, nutrient deprivation, and reactive oxygen species(1, 2). These stress conditions damage proteins in the cells by altering their functional conformations. The presence of unfolded and misfolded proteins disrupts proteostasis and can lead to cell death(3). The rapid induction of expression of a group of survival molecular chaperones known as heat shock proteins (Hsps) is critical for cytoprotective and protein structural repair purposes(2). This survival mechanism, known as heat shock response (HSR), is conserved among all organisms. Hsps were initially classified in families based on their molecular weight and further categorized as constitutive or inducible based on their steady-state expression levels(4). Constitutive and inducible members of the Hsp70 family play a key role in preserving protein homeostasis (proteostasis) because they prevent protein aggregation, assist in refolding damaged proteins, and clear aberrant proteins by cooperating with degradation machinery - the ubiquitin-proteasome system and autophagy(5). Due to its essential role in maintaining proteostasis, the cells need to tailor the levels of the inducible HSP70 to the extent of the misfolded protein burden. The expression of HSP70 is tightly regulated. First, to induce high levels during stress through the potent upregulation of its transcription and preferential mRNA translational. Then, to avoid its unnecessary accumulation during recovery by a rapid transcription shut-off and augmented *HSP70* mRNA instability. This fast switch from induction to attenuation of the HSR is critical to promote cell functionality. The persistent expression of the inducible Hsp70 under non-stress conditions causes growth defects in Drosophila(6) and promotes cancer formation in mammalian cells(7).

It is well known that under stress conditions, the robust transcription of HSP70 is induced by the activation of heat shock factor 1 (HSF1), followed by its binding to the heat shock element (HSE) in the promoter of *HSP70* genes(1, 8–12). The newly synthesized *HSP70* mRNAs engage in translation even though cap-dependent translation initiation and elongation are repressed to prevent the load of misfolded nascent polypeptides. The binding of eIF4F to the cap is inhibited through the recruitment of eIF4E by eIF4E Binding Proteins (4EBP) (13–15). Additionally, translation initiation is further dampened by the phosphorylation of eIF2α, which inhibits the GDP-GTP exchange(2, 10, 16–18). The co-transcriptional processing of the *HSP70* mRNA during stress favors its translation via a cap-independent pathway(19, 20). This mechanism involves co-transcriptional modifications of its 5’UTR to be recognized by the translation initiation factor eIF3 and the translation elongation factor eEF1A1 (21–24). Although how these and other yet-to-be-discovered factors act to translate *HSP70* mRNA is not fully defined. The ongoing translation stabilizes the *HSP70* mRNA during HS. During recovery, cells resume cap-dependent translation, and the *de-novo* synthesized HSP70 binds to the transactivation domain of HSF1 to self-repress its expression(2, 10, 11). This negative feedback loop between HSP70 and HSF1 reduces the HSR during recovery from stress(25). An additional pathway that contributes to the rapid shut down of HSP70 expression includes the efficient degradation of *HSP70* mRNA in a highly regulated fashion that requires the 3’ untranslated region (3’ UTR) for the instability of mRNA under permissive temperatures(26, 27). Therefore, *HSP70* mRNA switches from being a highly stable mRNA during stress to highly unstable during recovery(26, 27). Although the translation of *HSP70* mRNA is needed for its turnover, the factors tunning its fate in the cytoplasm in response to cellular stress status are unknown(10, 28).

The regulation of *HSP70* mRNA translation and stability relates to cellular changes in protein synthesis. The traditional translation model suggests that highly efficient translation initiation can increase the stability of mRNAs because more ribosomes are involved in the translation process, and they efficiently protect mRNA from degradation(29–35). Contrary to this model, it was recently found that a higher translation initiation rate can decrease protein expression from mRNAs containing pro-stalling codons in the budding yeast *Saccharomyces cerevisiae* (*S. cerevisiae*)(36–38). The increase in ribosome loading favors ribosome collisions between stalled ribosomes, which result in the formation of di-ribosomes (disomes) consisting of the leading ribosome and the subsequent colliding ribosome(39, 40). Colliding ribosomes signal to the ribosome quality control mechanism (RQC) to recycle stalled ribosomes and the mRNA surveillance mechanism – “No-Go Decay” (NGD) to trigger the degradation of the mRNA(31, 41, 42). The initiation of collision-associated quality-control mechanisms is mediated by Asc1 and Hel2 proteins (orthologous of RACK1 and ZNF598 in mammals, respectively), which stabilize the disomes(40, 42–46). Asc1 is a scaffold protein located at the head of the 40S subunit near the mRNA exit channel(47). In the context of RQC, the Asc1-Asc1 interaction between 40S subunits in a disome serves to stabilize the collision and provides an interface for recognition by the E3 ubiquitin ligases Hel2(40, 45, 48). Hel2 ubiquitinates the 40S ribosomal proteins eS10 and uS10 and promotes the splitting of stalled ribosomes(43, 46). Stabilization of collisions by Asc1 and Hel2 is necessary to recruit endonucleases that target the mRNA for degradation by NGD (49–51). Recent findings suggest that ribosome collisions and RQC do not necessarily trigger endonucleolytic cleavage and subsequent mRNA degradation by the NGD mechanism(49, 52–54). Ribosome stalling also signals to repress translation initiation to prevent the accumulation of partially synthesized peptides. In mammalian cells, translation is inhibited through ZNF598 recruiting GIGYF2 and 4EHP, which compete with eIF4E to bind to the cap of the mRNA undergoing ribosome stalling(55). *S. cerevisiae* lacks a 4EHP orthologue, but the absence of Hel2 triggers the phosphorylation of eIF2α to repress translation initiation through the recognition of ribosome collisions by the eIF2α kinase Gcn2 and the activation of the integrated stress response(53, 56).

Based on these studies, we hypothesized that the resumption of cap-dependent translation during recovery could link an increase in *HSP70* mRNA translation efficiency to its decay by NGD. To test this hypothesis, we used the yeast *S. cerevisiae* and studied the regulation of the four members of Stress Seventy sub-family A (Ssa), Ssa1-4(57). Ssa1p and Ssa2p are constitutively expressed, while Ssa3p and Ssa4p are inducible. *SSA3* mRNA translation initiation is regulated by an upstream open reading frame (uORF), leaving Ssa4p as the most inducible member of this subfamily. The codon sequence of *SSA4* mRNA is biased toward low-frequency codons, which can promote ribosome stalling and regulate Ssa4p expression by RQC and NGD. Accordingly, we found that RQC factors, Asc1 and Hel2, regulate *SSA4* mRNA life cycle in unexpected ways. Firstly, RQC downregulates Ssa4p synthesis during heat shock instead of recovery, preventing the overproduction of Ssa4p during stress. This regulation depends on the low codon optimality of the *SSA4* mRNA ORF and involves the ribosomal proteins Rps28A and Rps19B, which emerge as new RQC components. Secondly, RQC does not lead to NGD nor the degradation of *SSA4* mRNA during stress or recovery. Instead, Asc1p destabilizes *SSA4* mRNAs during recovery independently of its binding to the ribosome. This result points to two distinct functions of Asc1p that converge to control the life cycle of *SSA4* mRNA, and thus, we identified Asc1 as a major player in the regulation of the HSR in yeast.

## RESULTS

### The RQC factors Asc1p and Hel2p repress SSA4 mRNA translation during HS

Among the four members of the Ssa subfamily of yeast HSP70 proteins, the coding sequence (CDS) of *SSA4* mRNA is biased towards low-frequency codons when compared to the *SSA1*, *SSA2*, and *SSA3* mRNAs(58) (**Figure S1A**). The presence of non-optimal codons leads to slow decoding and stalling of ribosomes favoring ribosome collisions, especially in highly translated mRNAs (Park & Subramaniam, 2019). Therefore, we analyzed previously published ribosequencing data to identify the presence of stalled ribosomes over the CDS of the *SSA4* mRNA under heat-shock (HS) conditions (30 minutes (min) at 42°C) (59). The Pausepred analysis from the Trips-Viz program identified ribosomes stalled at position 400 (P1) on the *SSA4* mRNA on both experimental replicas and position 1800 (P2) in one of the replicas as peaks with 20-fold more ribosome occupancy than in the following position of the mRNA (60, 61) (**Figures 1A and S1B**). Although the codon sequence of *SSA4* mRNA protected by the ribosome was identical to two other *SSA* mRNAs, the *SSA4* mRNA was uniquely enriched in three low-frequency codons following the stalled ribosome. The presence of stalled ribosomes in the *SSA4* mRNA and its bias towards low-frequency codons led us to investigate the role of RQC and NGD mechanisms in regulating *SSA4* mRNA translation and decay, respectively (62).

**Figure 1.**
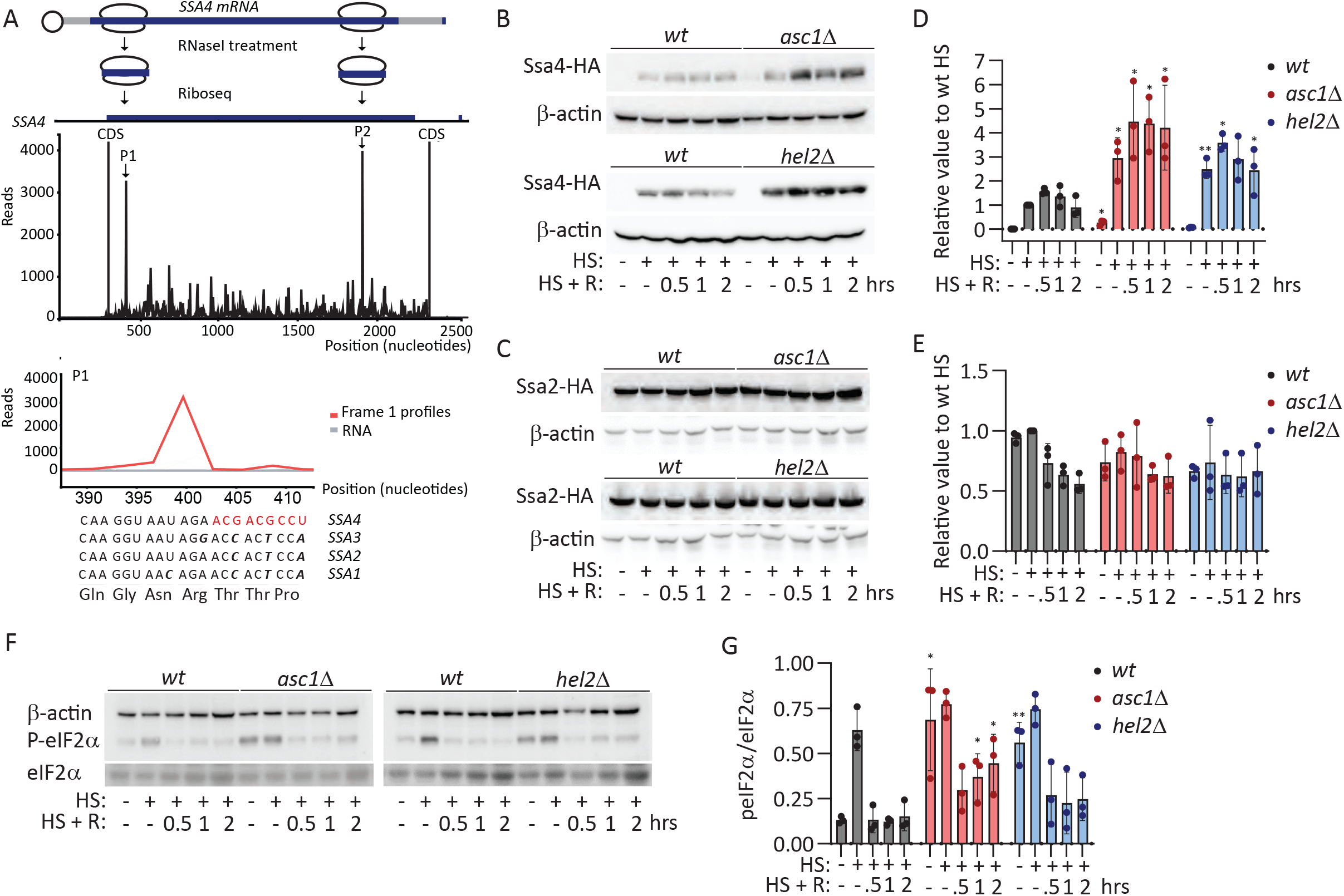
Depletion of Asc1p and Hel2p repress the translation SSA4 mRNA upon HS. **A**. Ribosome profiling analysis of *SSA4* mRNA in *Saccharomyces cerevisiae* upon 30 min of HS. Top: Schematic of *SSA4* mRNA with ribosomes stalled at two positions. Middle: Single transcript plot of ribosome sequencing analysis aligned with RNA-Seq data (Mühlhofer et al. 2019). This analysis found 2 major ribosome (P1 and P2 (indicated with black arrows)) stalled sites where the peak size for the Ribo-Seq data is 20x higher than the next peak. P2 was only detected in one of the replicas. Bottom: Magnification of the nucleotide sequence in covered by the ribosome at P1 and the encoded amino acids. **B-C.** Immunoblots to detect the expression of Ssa4p (**B**) and Ssa2p (**C**) tagged with 3 tandem HA epitopes and β-actin as the loading control in *wt* and *asc1Δ* and *hel2Δ* under basal [yeast growing at 25°C (HS: −) and (HS + R: −)], 30 min of HS at 42°C [(HS: +) and (HS + R: −)], and 30 min of HS at 42°C followed by half an hour (hr) (HS + R: 0.5), one (HS + R: 1), or two hrs (HS + R: 2) of recovery at 25°C. **D-E**. Quantification of the expression of Ssa4p and Ssa2p. The HA band intensity in Ssa4p (**D**) and Ssa2p (**E**) was first normalized to the β-actin band intensity for each condition and then related to the normalized expression of *wt* under HS. The bars indicate the mean and standard deviation (SD) of 3 independent experiments, each of them represented by a dot. N= 3, unpaired t-test (* = p<0.05, ** = p<0.001, *** = p<0.0001). **F.** Immunoblots to detect the phosphorylation of eIF2α. The expression of phosphorylated eIF2α (P-eIF2α), total eIF2α, and β-actin was quantified in *wt*, *asc1Δ*, and *hel2Δ* strains under the indicated basal, HS, and recovery conditions. **G**. Quantification of the phosphorylation status of eIF2α. The band intensity of P-eIF2α was divided by the total eIF2α for each condition. The bars indicate the mean and SD of 3 independent experiments, each represented by a dot. N= 3, unpaired t-test (* = p<0.05, ** = p<0.001).

The RQC factors, Asc1p and Hel2p, stabilize ribosome collisions and repress the translation of mRNAs with collided ribosomes(39, 41). To investigate the role of RQC factors in regulating Ssa4p synthesis during HS and subsequent recovery periods at permissive temperature, we depleted the *ASC1* or *HEL2* genes in haploid BY4741 wild type (*wt*) *S. cerevisiae*. Given the high similarity among the four *SSA* genes, we inserted 3x-Hemagglutinin (HA) epitopes in the C-terminus and 12xMS2V6 RNA stem-loops in the 3’UTR of each of the endogenous *SSA* genes to distinguish their proteins by immunoblot and mRNAs by single-molecule fluorescent *in-situ* hybridization (smFISH), respectively. Ssa1p, Ssa2p, Ssa3p, and Ssa4p were analyzed after HS at 42°C for 30 min and following 30, 60, and 120 min of recovery at 25°C in *wt*, *asc1Δ*, and *hel2Δ* strains. The intensity of the HA signal was normalized to the housekeeping β-actin, and the normalized value referred to the HS induction of the *wt* strain. The *asc1Δ* and *hel2Δ* strains induced a significantly higher Ssa4p expression during HS than the *wt* but did not enhance the expression of Ssa1p, Ssa2p, and Ssa3p nor the non-HS protein Doa1p during basal (25°C), HS, and recovery conditions (**Figures 1B-E, S1C and S1D**). The high expression levels of Ssa4p during HS persisted during the recovery, but the difference between wt and the *asc1Δ* and *hel2Δ* strains did not further increase.

We pondered two non-exclusive options to explain this result; Asc1p and Hel2p only repress *SSA4* mRNA translation during HS and/or the fast *SSA4* mRNA degradation at permissive temperature prevent Ssa4p synthesis during recovery. To investigate the first option, we considered that HS triggers the phosphorylation of eIF2α, favoring the translation of inducible *HSP* mRNAs and that in *asc1Δ* and *hel2Δ* cells, the Gcn2 kinase phosphorylates eIF2α even under basal conditions (53). Accordingly, *asc1Δ* and *hel2Δ* strains had higher eIF2α phosphorylation than the *wt* strain under basal conditions, and all strains phosphorylated eIF2α during HS. Interestingly, the phosphorylation of eIF2α quickly decreased to *wt* basal levels at 15 min of recovery even in the *asc1Δ* and *hel2Δ* cells, which can hinder *SSA4* mRNA translation during recovery and thus decrease the regulatory effect of Asc1p and Hel2p (**Figures 1F and 1G**). Overall, our results point to specific repression of *SSA4* mRNA translation by Asc1p and Hel2p during HS.

### Cells depleted of *ASC1* stabilized *SSA4* mRNA longer than *hel2Δ* and *wt* strains during recovery

Ribosome collisions can lead to endonucleolytic cleavage and the rapid decay of the problematic mRNAs by NGD (41, 63). Thus, we considered that ribosome collisions stabilized by Asc1p and Hel2p could lead to the rapid decay of *SSA4* mRNAs during recovery. We assessed the stability of *SSA* mRNAs by quantifying their expression levels at basal, HS, and several recovery times (15-, 30-, 60-, and 90-min) at 25°C after HS. We detected *SSA4, SSA3, SSA2, and SSA1* full-length mRNAs by Northern blot in the untagged strains using antisense probes that hybridized to their 3’UTR, which holds the most distinct nucleotide sequence among *SSA* genes. As expected, the expression of all *SSA* mRNAs was highly induced upon HS and rapidly returned to basal levels during recovery in the *wt* strain. The *asc1Δ* and *hel2Δ* strains only prolonged *SSA4* mRNA expression during recovery without affecting the decay of the rest of the *SSA* mRNAs (**Figures 2A, 2B, S2A, and S2B**). We plotted the intensity of the specific *SSA4* and *SSA2* mRNA bands as a control to calculate their half-lives by the non-linear regression that best fitted the curve connecting mRNA intensities during HS and recovery conditions. The effect on mRNA stability was specific to *SSA4* mRNA, for which *ASC1* depletion increased the half-life 2.5 times and *HEL2* depletion to 1.35 times (**Figure 2B**).

**Figure 2.**
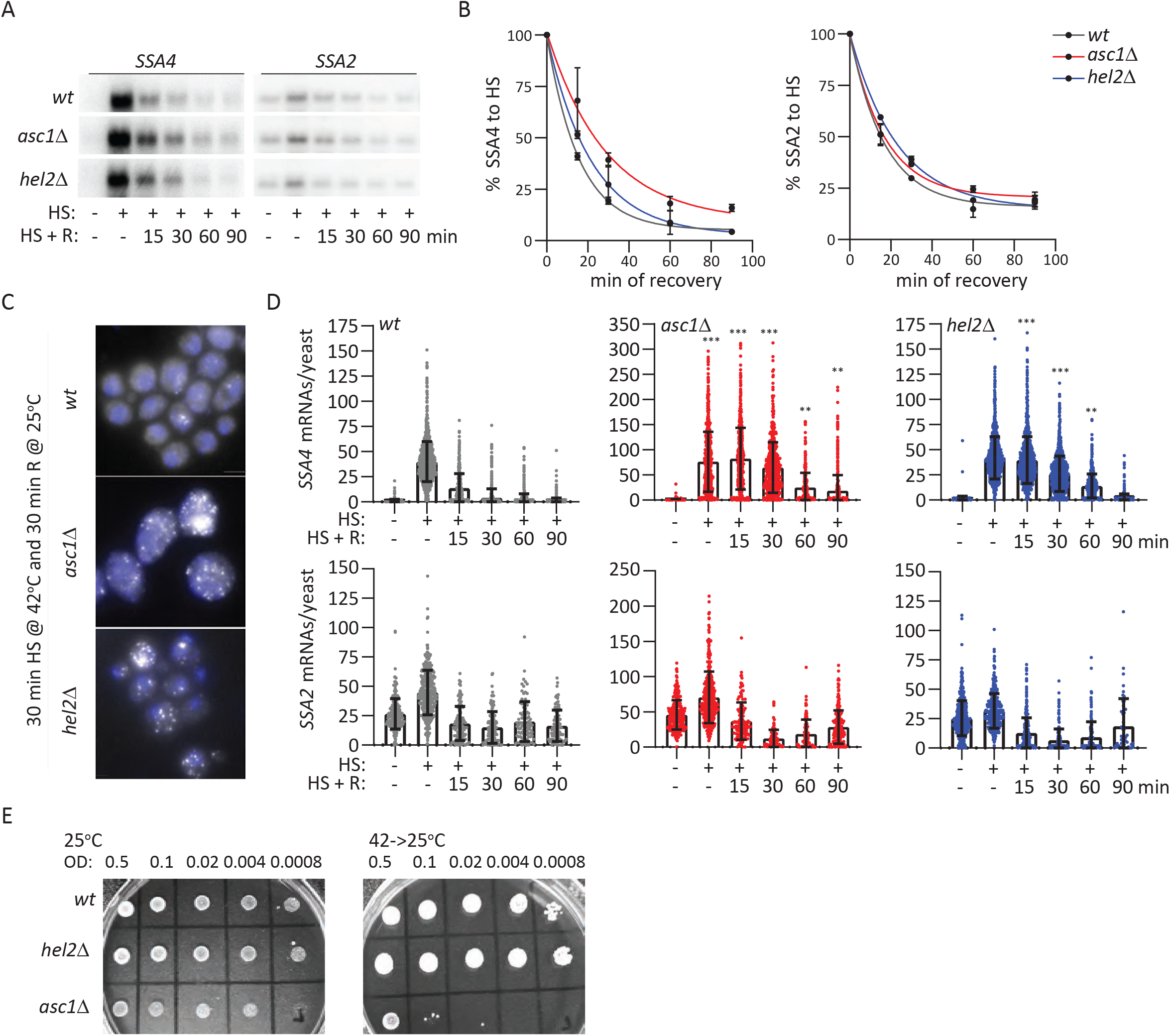
*SSA4* mRNA is stabilized in *asc1Δ* and *hel2Δ* cells during recovery. **A**. Northern blots to detect the expression of *SSA4* and *SSA2* mRNAs in *wt*, *asc1Δ*, and *hel2Δ* strains under basal [(HS: −) and (HS + R: −)], 30 min of HS at 42°C [(HS: +) and (HS + R: −)], and 30 min of HS at 42°C followed by 15 (HS + R: 15), 30 (HS + R: 30), 60 (HS + R: 60), or 90 (HS + R: 60) min of recovery at 25°C. **B**. Quantification of the half-life of *SSA* mRNAs during recovery. The band intensity of *SSA4* and *SSA2* mRNAs was normalized to the RNA loading by the methylene blue staining. Then, each recovery time was related to the intensity of the HS band (considered as 100% of induction) for each strain to obtain the decay curve and calculate the half-life. Dots of the curve indicate the average and SD of two independent experiments. **C**. Representative smFISH image to detect the expression of *SSA4-MS2V6* mRNA at 30 min recovery in *wt*, *asc1Δ*, and *hel2Δ* yeast strains. Scale bar 5 µm. **D**. Quantification of the number of *SSA4* or *SSA2* mRNAs per yeast as detected by smFISH. Bars indicate the average and SD of 3 experiments; each dot is the value of an individual yeast ((n=600-1200), t-test). **E**. Spot assay of *wt*, *asc1Δ*, and *hel2Δ* strains growth at 25°C and recovering at 25°C after 16 hrs incubation at 42°C. Numbers indicate the serial dilution.

We confirmed the prolonged expression of *SSA4* mRNA, not *SSA2* mRNA, in the *asc1Δ* and *hel2Δ* strains during recovery by smFISH in the *SSA4*- and *SSA2*-3xHA-12MSV6 tagged strains, respectively (**Figure 2C and 2D**). mRNAs were recognized with fluorescently labeled probes that hybridized to the MS2V6 sequence and quantified with the computational framework Fishquant (64). The average number of single *SSA4* mRNAs per cell is double in the *asc1Δ* strain than in the *wt* and *hel2Δ* strains, probably because they are larger cells and produce more mRNAs to compensate for their volume, as the *SSA2* was also more abundant (**Figure 2D**). In the *wt* strain, most *SSA4* mRNAs were cleared by 30 min recovery, while some *hel2Δ* cells and the majority of *asc1Δ* cells retained significantly higher numbers (*p* <0.05, t-test) of *SSA4* mRNAs until 90 min recovery. No significant differences among strains were found in the induction and decay of *SSA2* mRNA expression during HS and recovery (**Figure 2D**). By smFISH detection, the peak of *SSA4* mRNA expression was prolonged from HS to 15 min recovery in the *asc1Δ* and *hel2 Δ* strains. Since we could not quantify the contribution of nascent transcripts to the total mRNA pool, we attributed this difference with the Northern blot to the turnover of the cytoplasmic mRNA population in the first 15 min of recovery. The decay of *SSA4* mRNAs might be faster than the export of nascent transcripts to the cytoplasm in the *wt* cells but not in the *Asc1Δ* and *Hel2Δ* strains, and thus, they shifted the peak of expression to 15 min recovery.

Of the three strains, *asc1Δ* has the highest Ssa4 expression; thus, it should be better equipped to survive HS than *wt* and *hel2Δ* strains. However, as previously reported, the *asc1Δ* strain grew the slowest and was the most sensitive to HS, suggesting that its growth phenotype is independent of the expression of cytoprotective Ssa4p (**Figure 2E**). Overall, our results indicated that *asc1Δ* cells prolonged the stability of *SSA4* mRNA longer than *hel2Δ* cells, and therefore, Asc1p might play an additional role in the decay of *SSA4* mRNA during recovery from HS. Since the prolonged expression of *SSA4* mRNA during recovery did not further increase Ssa4p levels, Asc1p and Hel2p might regulate *SSA4* mRNA translation during HS and stability during recovery by independent mechanisms.

### Optimizing the *SSA4* coding sequence escapes RQC, but its mRNA is still stabilized in *asc1Δ* cells

To gain insight into the mechanisms behind Asc1p and Hel2p regulation of *SSA4* mRNA metabolism, we optimized the *SSA4* CDS to bypass ribosome stalling. We synonymized the *SSA4* CDS using a computational pipeline that utilizes the systems-level information and the codon context for the conditions under which genes are being expressed (65). Interestingly, the optimal *SSA4* CDS acquired the preferred codon usage of *SSA3*, *SSA2*, and *SSA1* mRNAs within their conserved amino acids, further supporting a specific bias in the *SSA4* CDS sequence towards low-frequent codons (**Figure S3**). To generate the *SSA4*-optimized (Opt) strain, we substituted the endogenous *SSA4* gene with the *SSA4-Opt* gene, which conserved the 5’- and 3’-UTR sequences but has a different CDS. Then we tagged the endogenous *SSA4*-*Opt* gene with 3xHA-12MSV6 to study the role of the *SSA4*-*Opt* CDS sequence in *SSA4* mRNA translation in *wt* cells (*Opt-wt*) and Asc1p and Hel2p roles in its translation regulation.

We found that *Opt-wt* cells expressed Ssa4p even in basal conditions, indicating that the biased of *SSA4*-*wt* CDS toward low-frequents codons prevent the spurious accumulation of Ssa4p in the absence of stress. Optimization of the *SSA4* mRNA sequence dramatically increased the upregulation of Ssa4p to twenty and forty-fold during HS and recovery, respectively (**Figures 3A and 3B**). Therefore, *SSA4-wt* CDS suppresses Ssa4p synthesis during HS. Remarkably, the induction of Ssa4p upon HS was similar in the *Opt-wt* and *Opt-hel2Δ* strains, which further demonstrated that Hel2p repression of Ssa4p synthesis depends on the low-frequent codons of *SSA4*-*wt* CDS (**Figures 3C and 3D**). Likewise, optimization of *SSA4* CDS decreased Ssa4p induction from 6-fold in the *wt-asc1Δ* to less than 2-fold in the *Opt-asc1Δ* strain during HS (**Figures 1B, 1D, 3C, and 3D**). We concluded that the non-optimal *SSA4-wt* CDS represses the translation during HS through the actions of Asc1p and Hel2p. We next investigated if *SSA4-Opt* CDS stabilizes *SSA4* mRNAs and prevents Asc1p and Hel2p from destabilizing them during recovery. We compared the expression and stability of *SSA4*-*Opt* mRNA and *SSA2* mRNA as a control in the *Opt-wt*, *Opt-asc1Δ*, and *Opt-hel2Δ* strains at 30 min HS at 42°C followed by 15-, 30-, 60- and 90-mins recovery at 25°C by Northern blot. Optimizing *SSA4* CDS made *SSA4* mRNA 1.5 times more stable than *SSA4-wt* during recovery and completely abolished the effect of Hel2p on *SSA4* mRNA stability. Therefore, increasing the translational efficiency of *Opt-SSA4* mRNA slightly increased its mRNA stability. Remarkably, the *Opt-asc1Δ* cells still extended 2.5 times the *SSA4* mRNA half-life without affecting the *SSA2* mRNA half-life (**Figures 3E and 3F**). This result supported the role of Asc1p in favoring the decay of *SSA4* mRNA during recovery, that it is independent of *SSA4* CDS and, thus, the NGD mechanism.

**Figure 3.**
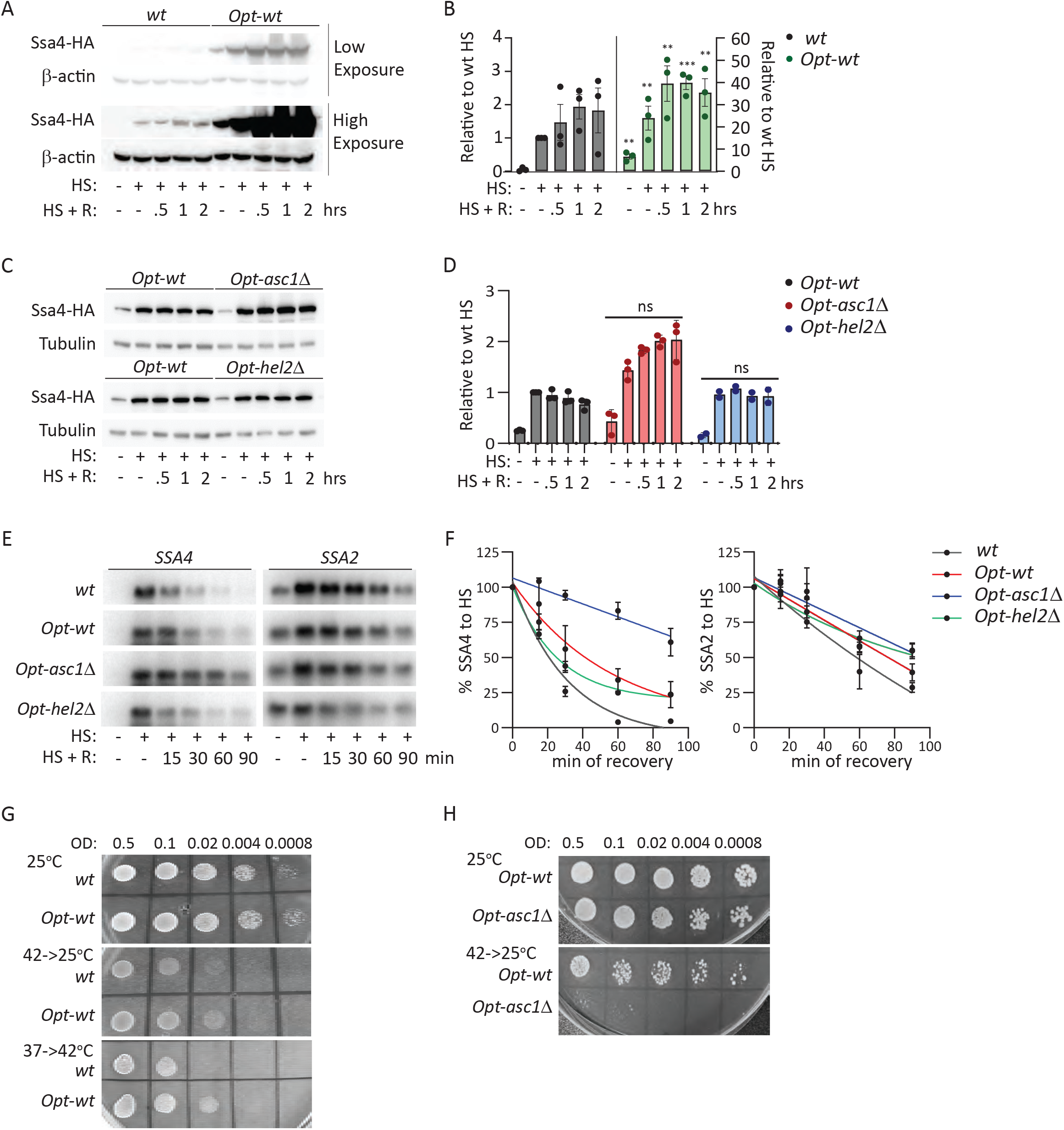
The optimized SSA4 mRNA escapes RQC preserves but is still destabilized by Asc1p. **A.** Immunoblots to detect Ssa4p-HA and β-actin as loading control in *wt* and *Opt* strains under basal, 30 min of HS at 42°C HS, and indicated recovery times. **B.** Quantification of the expression of Ssa4. The Ssa4-HA band intensity was first normalized to the β-actin band intensity for each condition and then related to the normalized expression of *SSA4-wt* yeast under HS. The bars indicate the mean and SD of 3 independent experiments, each represented by a dot. Unpaired t-test (* = *p*<0.05, ** = *p*<0.001, and *** = *p*<0.0001). **C.** Immunoblots to detect the expression of Ssa4p tagged with HA and Tubulin as loading control in *wt*, *Opt-wt*, *Opt-asc1Δ*, and *Opt-hel2Δ* under basal, 30 min of HS at 42°C HS, and indicated times of recovery. **D.** Quantification of the expression of Ssa4p. The HA band intensity in ssa4 tagged strains was first normalized to the tubulin band intensity for each condition and then related to the normalized expression of *wt* yeast under HS. The bars indicate the mean and SD of 3 independent experiments, each represented by a dot. Unpaired t-test (ns = non-significant, * = *p*<0.05, ** = *p*<0.001, and *** = *p*<0.0001). **E.** Northern blots detect the expression of *SSA4* (left) and *SSA2* (right) mRNAs in *wt*, *Opt-wt*, *Opt-asc1Δ*, and *Opt-hel2Δ* strains under basal, 30 min of HS at 42°C HS, and indicated times of recovery. **F.** Quantifying the half-life of *SSA4* (left) and *SSA2* (right) mRNAs during recovery. The band intensity of *SSA4* and *SSA2* mRNA was normalized to the RNA loading by the methylene blue staining. Then each recovery time was related to the intensity of the HS (considered as 100% of induction) for each strain to obtain the decay curve and calculate the half-life (t1/2 of *SSA4 OPT* mRNA: in *wt 22’*, *Opt-wt 40’*, *Opt-asc1Δ 100’*, and *Opt-hel2Δ 27’;* t1/2 of *SSA2* mRNA: in *wt 57’*, *Opt-wt 77’*, *Opt-asc1Δ 92’*, and *Opt-hel2Δ 118’*). **G and H**. Spot assay of *SSA4-wt* and *-Opt* (**G**) and *Opt-wt*, *Opt-asc1Δ*, and *Opt-hel2Δ* (**H**) strains growth at 25°C and recovering at 25°C after 16 hrs incubation at 42°C (42->25) and preconditioned by mild stress (37°C for 1 hr, 6 hrs at 25°C, and then incubated at 42°C, bottom). Numbers indicate the serial dilution.

Given the existing notion that HS maximized the production of Ssa4 to cope with protein misfolding, it was unexpected to find that *SSA4* CDS attenuates its own translation by the RQC mechanism. Thus, we investigated if Ssa4p overexpression could be toxic. The growth of the *wt* and *Opt-wt* strains was similar at the permissive temperature and during recovery from stress, and the overexpression of Ssa4p enhanced the survival of *Opt-wt* to HS upon preconditioning (**Figure 3G**). Overexpression of Ssa4p did not overcome the increased vulnerability of the *asc1Δ* cells to HS (**Figure 3H**). Thus, their inability to survive HS is independent of the role of Asc1p in regulating Ssa4p expression. Overall, our results support two separate functions of Asc1p in controlling the life cycle of *SSA4* mRNA; translational repression during HS, which is shared with Hel2p and depends on the *SSA4-wt* CDS, and mRNA decay during recovery that is independent of the CDS and *SSA4* mRNA translation efficiency.

### The coding sequence of the ASC1 locus but not the U24 snoRNA intron regulates SSA4 mRNA

In the *asc1*Δ strains, we attributed the regulation of *SSA4* mRNA translation and decay to Asc1p. However, the *ASC1* locus has an intron that leads to the expression of the small nucleolar RNA *snoR24*, known as *U24* (66, 67). *U24* is a C/D box snoRNA that guides 2'-O-methylation, also known as pseudouridylation of the 25S ribosomal RNA (rRNA), for which it requires at least 10 nucleotides (nts) of perfect complementarity (68–71). Sequence analysis of the *SSA4* 3’UTR revealed 10 nts of perfect complementarity with the *U24* sequence (TGAAGTAGCA) (**Figure 4A**). Since the 3’UTR sequence of *SSA4* mRNAs is required for the destabilization during recovery(27), we sought to determine whether the depletion of *U24* instead of Asc1p stabilizes *SSA4* mRNA in the *asc1Δ* strains. Thus, we restored either the expression of Asc1p or U24 from centromeric plasmids in the *asc1Δ* strain (as described in (66)). We also mutated the 3’UTR of the endogenous *SSA4* mRNA to prevent the binding of U24 (TG<u>*TTCAT*</u>GCA) and determine whether this sequence destabilizes *SSA4* mRNAs during recovery (*wt-3’UTR mut*). Northern blot analysis of the expression of *SSA4* mRNA in basal, HS, and recovery conditions showed that the sole expression of Asc1p destabilizes *SSA4* mRNA during recovery. On the contrary, the exogenous expression of *U24* in the *Asc1Δ* strain increased the stability of *SSA4* mRNA 3.4 times. Accordingly, mutating the *U24* binding sequence in the 3’UTR did not affect *SSA4* mRNA stability during recovery (**Figures 4B and 4C**). In order to validate our results, we investigated the role of Asc1p and *U24* in destabilizing *SSA4-Opt* mRNA. As expected, the expression of Asc1p was enough to destabilize *SSA4-Opt* mRNA in the *Opt-asc1Δ* and revert the half-life of *SSA4-Opt* mRNA to the *Opt-wt* levels. In contrast, *U24* did not change *SSA4-Opt* mRNA half-life and neither Asc1p nor *U24* changed *SSA2* mRNA stability in the *Opt-asc1Δ*. (**Figures 4B and 4C, and Figure S4B and S4C**). This result suggests that Asc1p regulated *SSA4* mRNA during recovery.

**Figure 4.**
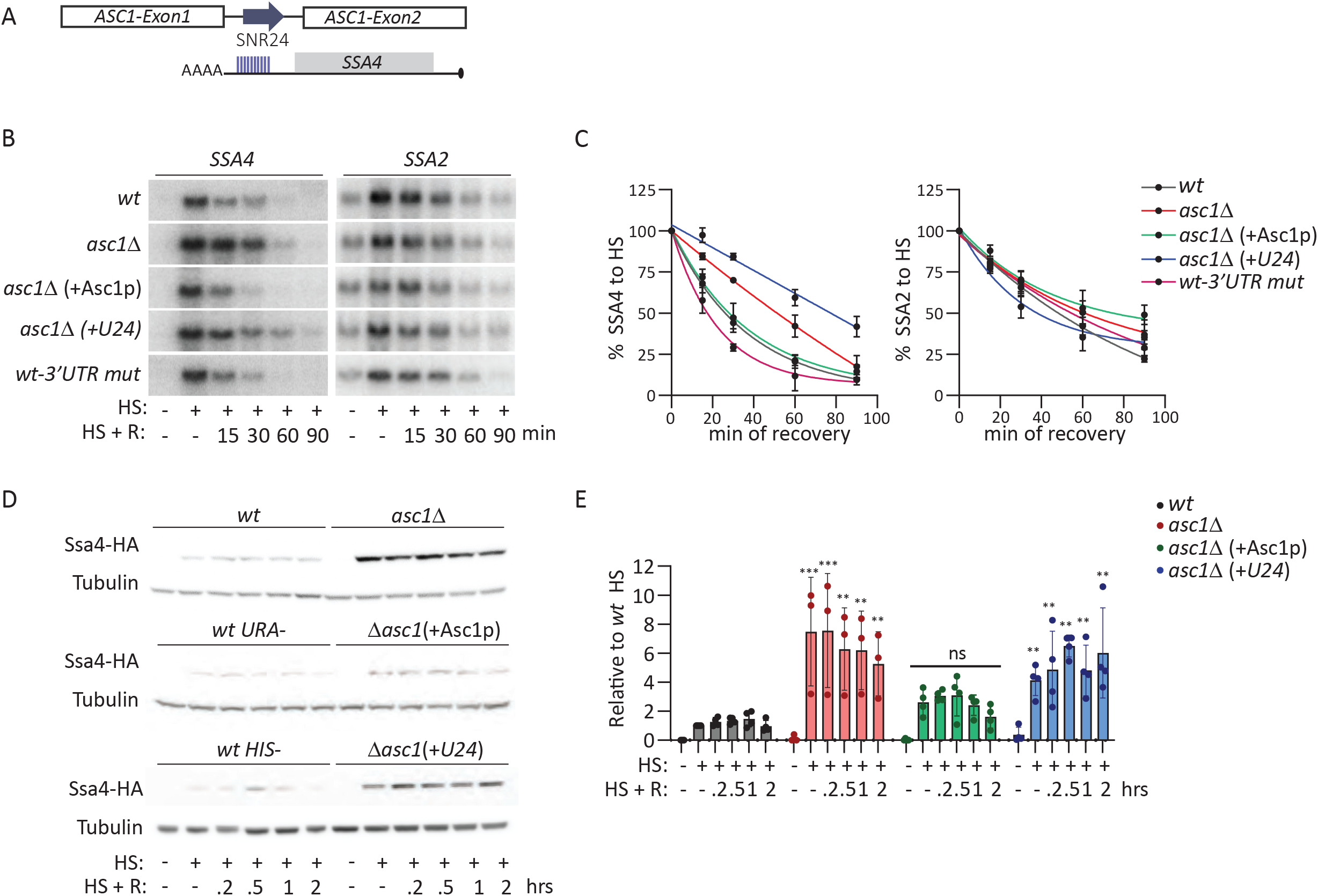
Asc1p, and not SNR24, regulates the SSA4 mRNA translation and stability. **A.** Schematic representation of *ASC1* gene locus containing the two exons and an intron, which leads to the expression of the small nucleolar RNA *SNR24* (*U24*) upon splicing. The 3’UTR of *SSA4* mRNA has a 10-nucleotide complementary region to *U24*, as indicated in blue lines. **B.** Northern blots to detect the expression of *SSA4* (left) and *SSA2* (right) mRNAs in *wt*, *asc1Δ*, *asc1Δ* expressing the CDS of Asc1p (*asc1Δ* +*Asc1p*), *asc1Δ* expressing only *U24 (asc1Δ*+*U24*), and *wt* strain with 5 mutations in the 3’ UTR of *SSA4* mRNA complement to *U24* (*wt-3’UTR mut*) under basal, 30 min of HS at 42°C HS, and the indicated recovery time points. **C.** Quantifying the half-life of *SSA4* (left) and *SSA2* (right) mRNAs during recovery. The band intensity of *SSA4* and *SSA2* mRNAs was normalized to the RNA loading by the methylene blue staining. Then, each recovery time was related to the intensity of the HS (considered as 100% of induction) band for each strain to obtain the decay curve and calculate the half-life (t1/2 of *SSA4* mRNA: in *wt* 28’, *asc1Δ* 51*’*, *asc1Δ* +*Asc1p* 21*’, asc1Δ* +*U24* 78*’, wt-3’UTR mut* 20*’;* t1/2 of *SSA2* mRNA: *wt* 48*’*, *asc1Δ* 60*’*, *asc1Δ* +*Asc1* 68*’, asc1Δ* +*U24* 36*’, wt-3’UTR mut* 55*’*). **D.** Immunoblots to detect the expression of Ssa4p tagged with three tandem HA epitopes and Tubulin as the loading control in *wt*, under basal, 30 min of HS at 42°C HS, and indicated recovery time points. **E.** Quantification of the expression of Ssa4p. The HA band intensity in *SSA4* tagged strains was first normalized to the tubulin band intensity for each condition and then related to the normalized expression of *wt* yeast under HS. The bars indicate the mean and standard deviation (SD) of three independent experiments, each represented by a dot. Unpaired t-test (** = *p*<0.001).

Although it is well known that Asc1p and Hel2p act together to regulate the translation of faulty mRNAs and trigger the RQC mechanism (46), we investigated whether *U24* also regulates *SSA4* mRNA translation. We quantified Ssa4p expression in the *asc1Δ* strains expressing either Asc1p or *U24*. While Asc1p expression restores the induction of Ssa4 to *wt* induction levels, yeast expressing *U24* in the absence of Asc1p sustained the higher Ssa4p expression of the *asc1Δ* strain (**Figures 4D and 4E**). Restoring either Asc1p or *U24* in the *Opt-Asc1Δ* did not change Ssa4p synthesis during HS and recovery. All together, these experiments indicate that U24 does not regulate the life cycle of *SSA4* mRNA and strongly support two independent roles of Asc1p on the regulation of *SSA4* mRNA life cycle; (1) mRNA translation during HS depends on low-frequent codons in the CDS, and (2) stability during recovery which is independent of the CDS and translation.

### Low binding of Asc1p to ribosomes destabilizes *SSA4* mRNA and partially regulates its translation

Asc1p is a multifunctional protein with roles in and out of the ribosome (67). We examined whether Asc1p binding to the ribosome is needed to destabilize *SSA4* mRNA during recovery and regulate its translation. We obtained three *ASC1* mutants, *M1X, DE, and DY*, as described by Thompson et al., 2016. The *M1X* mutant holds a substitution of the start by a stop codon in the *ASC1* CDS that prevents Asc1p expression but keeps *U24* levels. The *DE* mutant holds two substitutions, R38D and K40E, in the N-terminus that decrease Asc1p binding to ribosomes. The *DY* mutant, D109Y, has a lower ribosome binding capacity than the DE mutant and shows defects in NGD (67). Northern blot analysis to assess the effect of Asc1p null and ribosome-binding mutants *DE* and *DY* on *SSA4* mRNA stability revealed that only *M1X* cells prolonged the half-life of *SSA4* mRNA during recovery and none of the strains changed *SSA2* mRNA stability (**Figures 5A and 5B**). Firstly, this result confirmed our previous findings in the BY4741 background, showing that Asc1p expression, but not *U24*, destabilizes *SSA4* mRNA decay during recovery from HS. Secondly, since the *SSA4* mRNA half-life in the *DE* (24 min) and *D109Y* (27 min) were similar to the *wt* sigma strain (30 min), we concluded that the capacity of Asc1p to regulate *SSA4* mRNA stability is independent of its binding to the ribosome. These results were in line with Asc1p destabilizing *SSA4* mRNA independently of its CDS and codon optimality (**Figures 3E and 3F**).

**Figure 5.**
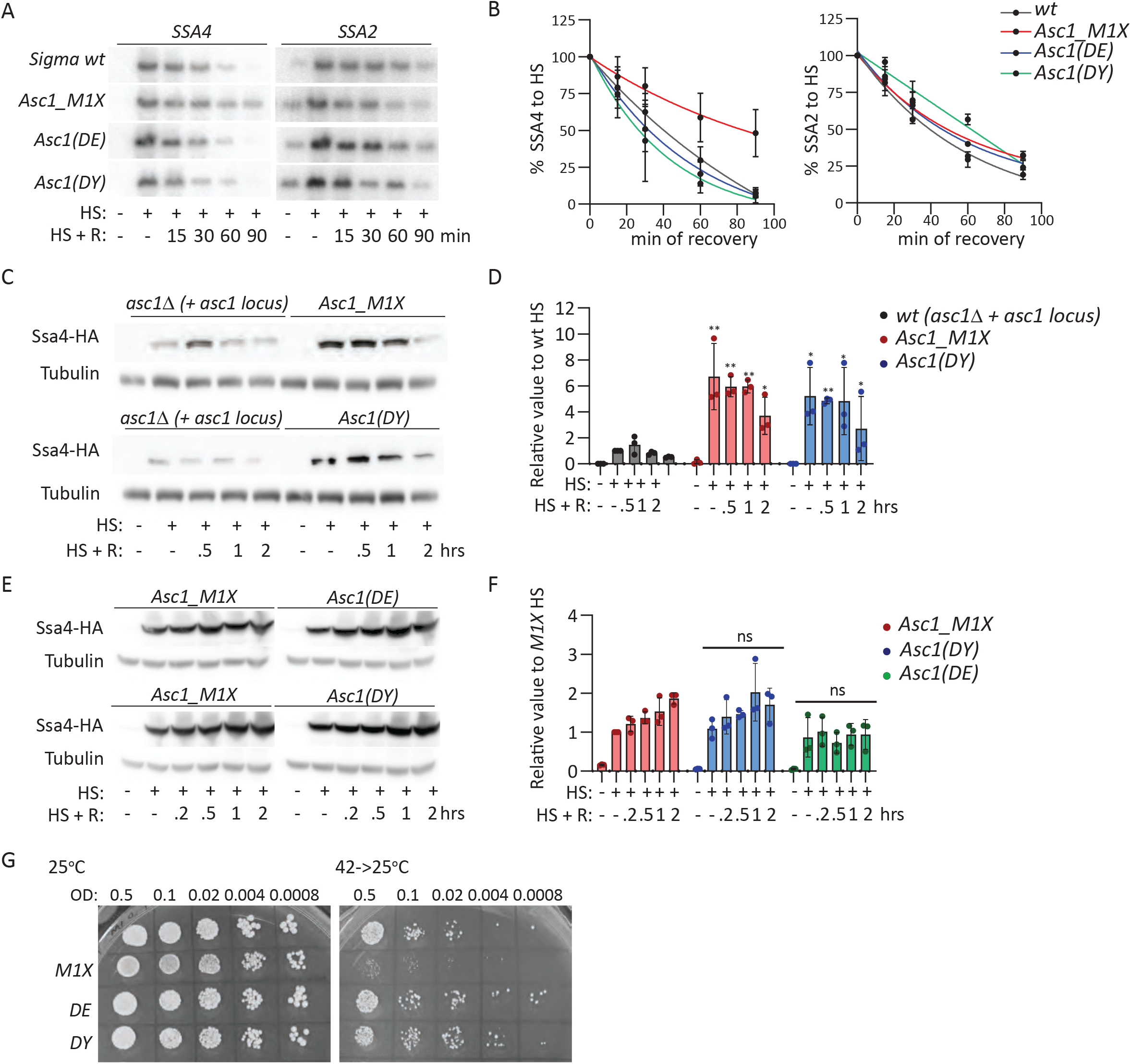
Role of Asc1p null and ribosome binding mutants on SSA4 mRNA stability and translation and survival to heat shock. **A**. Northern blots to detect the expression of SSA4 (left) and SSA2 (right) mRNAs in sigma *wt*, *Asc1_M1X*, *Asc1(DE)*, and *Asc1(DY)* under basal, 30 min of HS at 42°C HS, and indicated recovery time points. **B**. Quantify the half-life of *SSA4* (left) and *SSA2* (right) mRNAs during recovery. The band intensity of *SSA4* and *SSA2* mRNA was normalized to the RNA loading by the methylene blue staining, and then each recovery time was related to the intensity of the HS (considered as 100% of induction) band for each strain to obtain the decay curve and calculate the half-life (t1/2 of *SSA4* mRNA: in sigma *wt 30’*, *Asc1_M1X 84’*, *Asc1 DE* 24’and *Asc1 DY 27’;* t1/2 of *SSA2* mRNA: in sigma *wt 39’*, *Asc1_M1X 47’*, *Asc1 DE* 43’and *Asc1 DY 53’*). **C.** Immunoblots to detect the expression of Ssa4p tagged with three tandem HA epitopes and Tubulin as the loading control in *asc1*Δ BY4741 strains expressing full Asc1 locus (*wt*-phenotype), Asc1p mutant M1X and *DY* respectively under basal, 30 min of HS at 42°C HS, and indicated recovery time points. **D.** Quantification of the expression of Ssa4p. The HA band intensity in Ssa4 tagged strains was first normalized to the tubulin band intensity for each condition and then related to the normalized expression of *wt* yeast under HS. The bars indicate the mean and standard deviation (SD) of three independent experiments, each represented by a dot. Unpaired t-test (* = *p*<0.05 and ** = *p*<0.01). **E.** Immunoblots to detect the expression of Ssa4p tagged with three tandem HA epitopes and Tubulin as the loading control in sigma strains with Asc1p mutants *Asc1_M1X*, *Asc1(DE)* and *Asc1(DY)* respectively under basal, 30 min of HS at 42°C HS, and indicated recovery time points. **F.** Quantification of the expression of Ssa4p. The HA band intensity in Ssa4 tagged strains was first normalized to the tubulin band intensity for each condition and then related to the normalized expression of *M1X* yeast under HS. The bars indicate the mean and standard deviation (SD) of three independent experiments, each represented by a dot. Unpaired t-test, ns = no significant differences. **G.** Spot assay of Sigma *wt, M1X, DE* and *DY* under control (25°C, left), recovery (42°C for 16h, then incubated at 25°C, right) conditions plated on YPD. Numbers indicate the serial dilution.

We next investigated whether Asc1p should bind to the ribosome to repress Ssa4p synthesis. We used centromeric plasmids to express the *WT*, *M1X*, or *DY ASC1* genes in the *asc1*Δ BY4741 strain having the *SSA4* mRNA tagged with 3xHA-12MS2V6. Analysis of Ssa4-3XHA expression by Western blot revealed that the *M1X* strain expressed ~7 times more Ssa4p than the *WT* strain upon HS, as we previously found in the *asc1*Δ. Interestingly, the induction of Ssa4p in the *DY* strain is similar to that of the *M1X* strains upon HS, implying that the low binding of Asc1p to the ribosome is not enough to regulate *SSA4* mRNA translational repression (**Figures 5C and 5D**). These results were confirmed in the original sigma strains, whose expression of SSA4p was similar in the M1X, DE, and DY strains (**Figures 5E and 5F**). Therefore, binding of Asc1p to the ribosome is needed for the translational control of *SSA4* mRNA. We next investigated whether Asc1p must bind to ribosomes to promote the survival to HS. The expression of low ribosome-binding Asc1p mutants, *DE* and *DY*, provide *asc1*Δ yeast with the capacity to survive HS. Thus, the pro-survival role of Asc1p to HS is independent of its ribosomal binding and regulation of *SSA4* mRNA stability and translation (**Figure 5G**). Collectively, these results suggested that Asc1p repression of *SSA4* mRNA translation requires its partial binding to the ribosome, and Asc1p mediated destabilization of *SSA4* mRNA decay is independent of Asc1p binding to the ribosome. Thus, Asc1p probably uses two independent mechanisms to regulate SSA *4* mRNA translation and decay.

### HS enhanced the Asc1p binding to Rps28B and Rps19A, suppressing SSA4 mRNA translation

To identify the mechanisms and molecular partners sustaining the translation regulation and decay roles of Asc1p in *SSA4* mRNA metabolism, we Flag-tagged the endogenous Asc1p, immunoprecipitated it in basal and HS yeast and proceeded to liquid chromatography-mass spectrometry. HS did not induce any posttranslational modifications of Asc1p but significantly changed its interactome. HS enhanced the interaction of Asc1p with 12 proteins and decreased its interaction with 74 proteins (Fold change>2 (**Figures 6A and Table S4**)). Among the 13 proteins preferentially bound by Asc1p upon stress, six were chaperones (HSC82 and proteins labeled in red circles), suggesting that Asc1 unfolds during heat shock, and two were small ribosomal proteins. From the 13 proteins, the Ste20p kinase, the ribosomal protein Rps28Ap and Yra1p were of interest because they regulate the metabolism of mRNAs (72–74). Previous literature found that Asc1p phosphorylates Ste20p, which phosphorylates the decapping enzyme Dcp2p affecting the translation and decay of some mRNAs upon stress (73). Rps28A locates in the 40s ribosomal but also functions outside the ribosome to degrade *YRA1* pre-mRNA and *RPS28B* mRNA through the interaction with the enhancer of mRNA decapping protein 3 (Edc3p) (72). Hence, we depleted *STE20*, *RPS28A*, and *EDC3* genes to analyze their impact on *SSA4* mRNA translation. First, we studied the role of Ste20p, Rps28A, and Edc3p in destabilizing *SSA4* mRNA. The decay of *SSA4* and *SSA2* mRNAs were analyzed by Northern blot and showed that *ste20Δ* and *wt* had similar half-lives. In r *ps28AΔ* and *edc3Δ* cells, the half-life of *SSA4* mRNA was prolonged 1.5 times and did not affect *SSA2* mRNA stability. Nonetheless, the increase in *SSA4* mRNA stability was lower than the depletion of *ASC1.* We concluded that neither Ste20p, Edc3p nor Rps28Ap function with Asc1p outside the ribosome to destabilize *SSA4* mRNA during recovery. The *SSA4* mRNA degradation during recovery depends on Dhh1 and Xrn1, and it is independent of the Not4p and Ski7p, suggesting a role of *Asc1p* in activating factors that decap the *SSA4* mRNA and favor its degradation by the exonuclease Xrn1p from 5’ to 3’ (**Figures S5A and S5B**). It is important to note that the prolonged half-life of *SSA4* mRNA in *xrn1Δ* and *dhh1Δ strains* did not result in an increased upregulation of Ssa4p during recovery, which indicates that the translation of *SSA4* mRNA is suppressed during recovery (**Figures S5C and S5D**).

**Figure 6.**
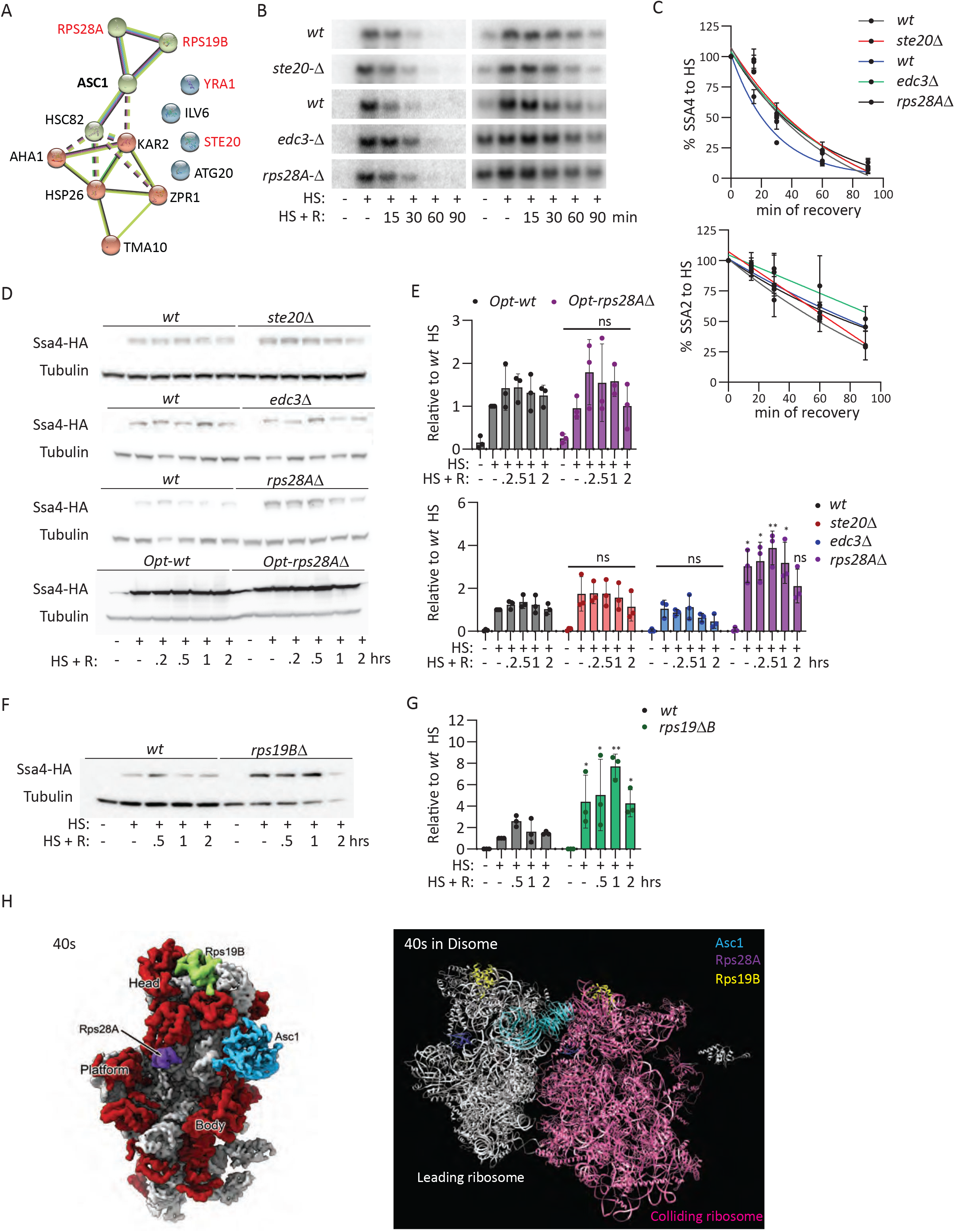
Ribosomal proteins Rps28Ap and Rps19B interact with Asc1p during HS to repress Ssa4p synthesis. (**A**) Asc1p interaction network of proteins significantly enriched (> 2-fold) at 30 min of HS plotted with STRING. (**B**) Northern blots to detect the expression of *SSA4* (left) and *SSA2* (right) mRNAs in *wt* vs *ste20Δ* (top) and *wt*, *edc3Δ* and *rps28AΔ* under basal, 30 min of HS at 42°C HS, and indicated recovery time points. (**C**) Quantification of the half-life of *SSA4* (left) and *SSA2* (right) mRNAs during recovery. The band intensity of *SSA4* and *SSA2* mRNA was normalized to the RNA loading by the methylene blue staining and then, each recovery time was related to the intensity of the HS (considered as 100% of induction) band for each strain to obtain the decay curve and calculate the half-life (t1/2 of *SSA4* mRNA: in *wt* 32’, *ste20Δ 35’, wt 22’*, *edc3Δ 35’*, 43’and *rps28AΔ 34’;* t1/2 of *SSA2* mRNA: in *wt 70’*, *ste20Δ 54’, wt 81’*, *edc3Δ 92’*, and *rps28AΔ 78’*). **D**. Immunoblots to detect the expression of Ssa4p tagged with three tandem HA epitopes and Tubulin as the loading control in *wt, ste20Δ*, *edc3Δ* and *rps28AΔ* yeast and *Opt-wt* and *Opt-rps28Δ* strains under basal, 30 min of HS at 42°C HS, and indicated recovery time points. **E.** Quantification of the expression of Ssa4p. The HA band intensity in Ssa4p tagged strains was first normalized to the tubulin band intensity for each condition and then related to the normalized expression of *wt* yeast under HS. The bars indicate the mean and standard deviation (SD) of three independent experiments, each represented by a dot. Unpaired t-test (ns = non-significant, * = *p*<0.05 and *p*<0.001). **F.** Immunoblots to detect the expression of Ssa4p tagged with three tandem HA epitopes and Tubulin as the loading control in *wt* and *rps19BΔ* yeast strains under basal, 30 min of HS at 42°C HS, and indicated recovery time points. **G.** Quantification of the expression of Ssa4p. The HA band intensity in Ssa4p tagged strains was first normalized to the tubulin band intensity for each condition and then related to the normalized expression of *wt* yeast under HS. The bars indicate the mean and standard deviation (SD) of three independent experiments, each represented by a dot. Unpaired t-test (* = *p*<0.05 and *p*<0.001). **H.** Structure of the *S. cerevisiae* 40s as a monomer (left, surface representation) and as a disome (left, ribbon representation) with colored Asc1p, Rps28Ap, and Rps19Bp.

Given the tighter interaction of Rps28A and Ste20p with Asc1p during heat shock, we next investigated their role in regulating *SSA4* mRNA translation. As expected, neither Ste20p nor Edc3 affects the synthesis of Ssa4p during heat shock and recovery. However, *rps28AΔ* cells synthesized five times more Ssa4p than *wt* cells. This induction is comparable to the one exhibited by *asc1Δ* cells (**Figures 1B, 1D, 4D and 4E**). To demonstrate that the novel role of Rps28A in regulating *SSA4* mRNA translation depends on its binding to the ribosome and the presence of low-frequent codons, we depleted *RPS28A* in the SSA4-Opt strain. In this case, the absence of Rps28A did not affect the expression of Ssa4p during HS, pointing to Rps28A as a new ribosomal component of the RQC mechanism (**Figures 6D and 6E**). Since <u>Rps19B</u> interaction with Asc1p is also enhanced during heat shock, we investigated its role in regulating *SSA4* mRNA translation during heat shock. Interestingly, *rps19BΔ* also induced significantly more Ssa4p than *wt* cells during heat shock (**Figures 6F and 6G**). Thus, Rps19Bp is also a novel ribosomal protein regulating the translation of *SSA4* mRNA during heat shock, probably through the RQC mechanism. We identified the position of Asc1, Rps28A, and Rps19B proteins in both the 40S subunit and in the published structure of the yeast disome (40), but we did not observe a direct interaction among them (**Figure 6H**). Since these structures have been obtained from yeast growing under permissive temperatures, the dynamic interaction of ribosomal proteins and their structure under stress is unknown. Therefore, we speculate the increased interaction of these ribosomal proteins upon heat stress could be either by HS-mediated alteration in the 40S ribosome structure and/or position or could be mediated by an additional factor(s). Taken together, our results further corroborate the role of the ribosome and *SSA4* CDS in regulating its translation during HS and discover Rps28A and Rps19B as ribosomal proteins inhibiting the translation of *SSA4* mRNA during heat shock.

## DISCUSSION

Cells rapidly adapt to survive harsh environmental conditions through the potent upregulation of Hsp’s expression. Members of the HSP70 family were initially detected as the most upregulated Hsps. The regulatory elements leading to such fast and transient activation were soon identified in the HSP70 promoter, the HSE to direct transcription, and the UTRs, the 5’UTR to mediate translation and the 3’UTR to regulate mRNA stability (10). Our work demonstrates that the coding sequence of the *SSA4* gene, the most induced HSP70 in yeast, also regulates its expression. Surprisingly, the bias of the CDS toward low-frequency codons lessens the extent of Ssa4p synthesis during heat shock by activating the RQC mechanism, which feedbacks to repress its own translation. Hence, our data argue that not all stress-induced gene expression pathways act to uplift HSP70 expression. Recently, a mechanism to attenuate HSP70 synthesis was found in mammalian cells during heat shock. In this case, the regulation relies on the heat-induced non-coding RNA *Heat* that ameliorates HSP70 transcriptional induction (75). In addition to the role of the *SSA4* mRNA CDS during heat shock, we also discovered that it is biased towards low-frequent codons and also prevents the spurious accumulation of Ssa4p under permissive growth conditions. This observation suggests that *SSA4-wt* CDS lessens the translation of the few *SSA4* mRNAs transcribed at permissive temperatures, providing an extra checkpoint to tailor HSP70 synthesis to the burden of misfolded proteins. Since the four Ssa proteins have a high amino acid identity (over 80%), the used of different codons offers the means for a different translational regulation.

In our study, we fully reverted the CDS of *SSA4* to optimal codons used in heat shock (65), and thus, codon optimality instead of stretches of polybasic amino acids cause ribosome stalling (31, 41, 76, 77). Most experiments designed to study the RQC and NGD components relied on reporters and were undertaken under permissive conditions. These reporters hold a stretch of polybasic or rare amino acids or focus on mRNAs regulated by the non-stop-decay (NSD) mechanism, which allows the translation of the polyA tails leading to stretches of the basic amino acid arginine(52, 78–80). We revealed *SSA4* as one of the few endogenous mRNAs whose translation is regulated by ribosome collisions. For *SSA4* regulation, the RQC mechanism is restricted to heat shock because neither the depletion of *ASC1* or *HEL2* nor its CDS optimization augmented Ssa4p accumulation during recovery. This result implies a mechanism boosting *SSA4* mRNA translation under heat shock, which probably depends on eIF2α phosphorylation because *asc1Δ* and *Hel2Δ* cells exhibited basal eIF2α phosphorylation only at early recovery time points. Identifying these factors will help elucidate the mechanism that autoregulates the expression of Ssa4p by blocking the initiation of *SSA4* mRNA translation in response to ribosome stalling in yeast. In mammalian cells, RQC signals to repress translation initiation by ZNF598 recruitment of GIGYF2 and 4EHP, which binds to the cap of the mRNA holding stalled ribosomes(55). Yeast cells do not have a 4EHP orthologue, and the two proteins with GYF domains, Syh1p and Smy2p, have not been related to translation regulation.

The RQC and NGD mechanisms are intimately connected in yeast(42, 81). Besides, recent work has shown that stall ribosomes can signal to the CCR4-NOT complex via Not5 to deadenylate the mRNA and trigger the degradation(49). However, NGD did not trigger *SSA4* mRNA decay, as shown by its high stability during heat shock and the discrete increase in the half-life of the optimized over the *wt SSA4* mRNA during recovery. It was unexpected to find Asc1p taken upon the role of destabilizing *SSA4* mRNA during recovery and doing this function independently of the CDS and its binding to ribosomes. Asc1p is a multifunctional protein with diverse roles in and out of the ribosome(67). We first rationalized that its capacity to phosphorylate the kinase Ste20, which regulates the decapping enzymes Dcp2 and Dhh1p, could lead to the degradation of *SSA4* mRNA by the exonuclease Xrn1(73, 82). Still, depletion of Ste20 did not stabilize *SSA4* mRNA. It might be possible that the mechanism used by Asc1p to destabilize *SSA4* mRNA is independent of direct interaction with a regulatory factor. Instead, Asc1p capacity to regulate the assembly of processing bodies (PBs) might facilitate the release of decay enzymes that degrade *SSA4* mRNAs(83). Since the formation of condensates is critical for cell survival to stress (84, 85), it is tempting to speculate that the role of Asc1 in assembly PBs might explain the inability of *asc1Δ* to recover from heat stress.

Our work establishes a new role for the RQC mechanism in regulating the expression of the inducible Ssa4p upon heat stress and defines the ribosomal proteins as new RQC components. Future experiments will determine if Rps28A and Rps19B only operate under stress conditions and they solely regulate *SSA4* mRNA translation or if they also mediate RQC under permissive growth conditions. Interestingly, the RQC essential factor Asc1p is the only one to mediate the decay of *SSA4* mRNA during recovery. It was unexpected to find that it uses a mechanism unrelated to the codon composition and Asc1p binding to the ribosome. Thus, Asc1p operates two synergistic mechanisms that converge to regulate the HSP70 expression during stress and recovery conditions. If these roles are conserved in mammalian cells, RACK1 might appear as a critical factor in tuning the outcome of the HSR and a putative therapeutic target to recover proteostasis under pathological conditions like cancer and neurodegeneration.

## MATERIALS AND METHODS

### Yeast culture

All yeast strains are derived from the parental strain BY4741, and their genotypes are summarized in **Supplementary Table 1**. They were grown in yeast extract peptone dextrose (YPD) media or the conditional media appropriate for their genotype at 25°C with constant shaking at 250 rpm. The knock-in and depleted strains were created by homologous recombination of the parental strain after transformation of the PCR fragment amplified from a plasmid carrying selection-specific markers, as previously described(86). Gene depletions and knock-ins were verified by PCR of genomic DNA extracted from individual colonies, as previously described. Primers and plasmids are listed in **Supplementary Tables 2 and 3**.

### Heat shock and recovery

For northern and western blot experiments, cells in the logarithmic growth phase (0.4-0.6 OD:600) were heat shocked at 42°C in a water bath with constant shaking at 80 rpm for the indicated collection time points. Immediately after HS the heated media was replaced by the same volume of media stored at room temperature (RT). The culture flasks were then placed back in a shaker incubator maintained at 25°C and rotated at 250 rpm until the collection and cultures were collected for downstream sample processing. For the spot assays, cells at 1.0-1.5 OD:600 were diluted in water to 0.5 OD:600 and then serially diluted in the ratio 1:5 in water five times. Five µl of the serially diluted cells were plated in YPD-agar plates, which were incubated at 25°C or HS in an incubator at 42°C for 16 hours (h) and then placed at 25°C. The precondition was done by HS at 37°C followed by 5 h of recovery at 25°C before moving the plates to 42°C. The plates were then checked for colonies every 24 hours and taken pictures.

### Protein extraction and Immunoblotting

Five ml of unstressed, heat-shocked, or yeast undergoing recovery were collected and centrifuged at 4000rpm for 5 min. The cell pellet was first washed with 2M LiOAc at RT and then with 0.4M NaOH on ice. Cells were lysed with SDS-PAGE buffer (60mM Tris-HCl pH 6.8, 10% glycerol, 2% SDS, 5% 2-mercaptoethanol, 0.0025% bromophenol blue). The lysates were heated for 10 min at 95°C before loading on a 10% SDS page gel and transferred to a nitrocellulose membrane. Ponceau S stain checked equal protein loading. The membrane was then blocked with 5% skim milk in PBST (1xPBS, 0.05% Tween 20) for 1 h at RT and then incubated with specific antibodies (eIF2α, P-eIF2α (Cat# 9722, 9721S, Cell Signaling™), HA (Cat#901501, BioLegend™), Tubulin (Developmental Studies Hybridoma Bank), β-actin (Cat# A2228, EMD Millipore Corp™) overnight at 4°C. Followed by 3 washes in PBST, the membrane was incubated with horseradish peroxidase-conjugated goat-anti-mouse (Cat#1706516, Biorad) or goat-anti-rabbit antibody (Cat#1706515, Biorad) for 2 h at RT. Three washes were performed, followed by ECL treatment and imaging with Chemidoc imaging system from Biorad. The intensity of the target protein signal was quantified using ImageJ software and normalized to that of loading control (β-actin or Tubulin).

### RNA extraction and Northern Blotting

Five ml of unstressed, heat shocked, or yeast undergoing recovery were pelleted by spinning at 4000 rpm for 5 min at 4°C. The pellets were resuspended in 0.5 ml of RNA extraction lysis buffer (10mM Tris-HCl pH8.5, 5mM EDTA, 2% SDS, 2% stock 2-mercaptoethanol), and the contents were transferred to 1.5 ml tubes. Cells were lysed by incubating the tubes in a heat block at 83°C for 20 min. After Spinning at 12000 g for 5 min, the supernatant was transferred to a fresh tube containing 0.55 ml of pH 8 phenol. After vortexing for 30 seconds, the top layer was transferred to a new set of tubes labeled N following a spin at 12000g for 2 min. To the previous tube, 0.25ml of the RNA extraction lysis buffer was added and vortexed briefly. An equal volume (0.25ml) of Chloroform was added, vortexed and spun to transfer the top layer to tube N. Another 0.55ml of pH 8 phenol was then added to N tubes, vortexed, spun, and top layer is once again transferred to a new tube containing 0.55ml of Acid Phenol-Chloroform, pH 4.5. The tubes were vortexed briefly, spun and 0.45ml of top layer was transferred to a new set of tubes containing 0.6M sodium acetate, pH 4.5. The contents were mixed by flicking followed by a quick spin. Once again, Acid phenol-Chloroform, 0.6ml was added to the tubes accompanied by a vortex and spin. Around 0.35ml of top layer was once again transferred to tubes with 1.1ml 100% Ethanol and 0.03ml of 5M Ammonium acetate. After being mixed, the samples were placed in −80°C overnight. Next day, the samples were spun at 4°C for 15 min and the supernatant was discarded. The pellet was washed twice with 80% ethanol and allowed to air dry. The dried pellet is dissolved in 0.04ml of RNase-free water and the RNA was quantified. Equal amount (1000-2000 ng) of RNA was aliquoted into fresh tubes and speed vac dried for 45 min at 45°C. The samples were then resuspended in 5µl of RNase free water and mixed with 7µl of homemade RNA loading dye. The RNA samples were run in a 0.75%-1% denaturing gel in 1X MESA buffer. The transfer to zeta probe nylon membrane was set up using capillary electrophoresis overnight. The membrane was UV crosslinked at 1200mJ, stained for total RNA, hybridized, and exposed to phosphorscreen, and developed using phosphorimager. The northern blotting and the radiolabelling of probes were performed as mentioned in (87). Genomic BY4741 was used as a template to PCR amplified probes that target SSA1, SSA2, SSA3, SSA4, SSA4-MS2, SSA4 OPT 3’ UTR, SSA4 OPT 5’UTR are provided in **Supplementary Table 2.**

### Calculating half-life of mRNAs

The northern blots were quantified using imageJ and corresponding methylene blue staining was quantified to normalize for the loading. Considering the intensity of HS (timepoint: 0) to be 100%, the relative intensity to HS was calculated for all the recovery samples (timepoint: 15, 30, 60, 90 min). A polynomial curve was plotted for time *vs* percentage of mRNA decayed. The polynomial equation was obtained for the curve and solved for X, given that Y is 50 using what-if analysis under the data tab in Microsoft Excel.

### Computing the percentage of preferred codons

A list of preferred codons in *S.cerevisiae* is procured from Bennetzen & Hall, 1982 and the coding sequence of SSA mRNAs is obtained from saccharomyces genome database (https://www.yeastgenome.org/). Using the python program (github link: https://github.com/LR-MVU/YEAST-SSA.git), the occurrence of each preferred codon was counted. Then the sum of each preferred codon is divided by the total number of codons to calculate the fraction of preferred codons. A bar chart was plotted in percentages of preferred codons in individual SSA mRNAs in *S.cerevisiae*.

### Single-Molecule in situ hybridization (smFISH) and Imaging analysis

The smFISH procedure was done as previously described with no modifications (88). Briefly, yeast strains were grown in 25 ml at 25°C to early log phase in YPD and heat shocked. At the indicated time point, they were fixed in 4% paraformaldehyde, permeabilized in spheroplast buffer containing lyticase, and seeded onto poly-L-lysine coated coverslips. After ethanol incubation, rehydration with 2X SSC, and prehybridization, they were hybridized with Stellaris smFISH probes to detect MS2V6 or MDN1 (LGC Biosearch Technologies) as previously designed (Tutucci et al, 2018). The coverslips were washed, dried, and mounted in Prolong gold antifade mounting media (Invitrogen). The smFISH experiments were imaged using a wide-field inverted Nikon Ti-2 wide-field microscope equipped with Spectra X LED light engine (Lumencor), and Orca-Fusion scMOS camera (Hamamatsu) controlled by NIS-Elements Imaging Software. For yeast cells, a 100x 1.49 NA oil immersion objective lens (Nikon) was used with xy pixel size 67.5 nm and z-step 200nm. The outlines were created by CellProfiler pipeline, and the single mRNAs were quantified using FISHquant software (Mueller et al, 2013).

### Ribosome profiling analysis

Ribosome profiling and RNA seq data in yeast under heat shock conditions from Mühlhofer et al., 2019 (Cell Reports 2019) were downloaded from GEO datasets. The accession numbers of Riboseq data: SRR9265440 and SRR9265438. The accession numbers of RNA seq data: SRR9265437 and SRR9265428. Raw sequencing reads were processed first by trimming adapters using Cutadapt 3.4 and discarding the low-quality reads; before alignment to yeas transcriptome, the reads were mapped to rRNA, tRNA and aligned reads were discarded. The remaining reads were then mapped to yeast transcriptome and the resulting SAM files were further processed to SQLite file using a python script from Trips-Viz webserver (https://trips.ucc.ie/) for compatibility with the server. PausePred (61), the built-in function of Trips-Viz (60) with the default setting, was used to find the ribosome pausing site. The pausing site for SSA4 mRNAs was visualized using Trips-Viz.

### Protein IP and Mass spectrometry

Flag tagged Asc1 (Asc1p-3xFlag) yeast cultures were grown in 400 ml at 25°C to early log phase in YPD and heat shocked for 1 hour at 42°C. The cells were then pelleted by centrifugation 3750rpm for 3 minutes in precooled centrifuge. The pellets were washed with 10ml water and aliquoted to eppendorf tubes. The pellets were then snap frozen with liquid nitrogen in and then thawed on ice. The pellets were then resuspended in 2ml of lysis buffer (100mM HEPES pH 8.0, 20mM Magnesium Acetate, 10% Glycerol, 10mM EGTA, 0.1mM EDTA) containing protease and phosphatase inhibitors. Glass beads (0.5mm) were then added and vortexed for 20 times with 30 s on-off cycles in the cold room. The lysate was separated from the beads by centrifugation and transferred the crude extract to a new tube. Alongside, the M2-anti-Flag beads by prepared by washing 3 times with 500ul of lysis buffer. The crude extract was then incubated with the beads on a nutator for 2 hours in a cold room. The beads were subjected to a magnetic field and the flowthrough was collected. The beads were then washed thrice with 1ml of lysis buffer and each wash was collected. The samples were run on SDS-PAGE gel including the beads, crude extract, flow through, washes and protein standards in SDS-PAGE buffer (60mM Tris-HCl pH 6.8, 10% glycerol, 2% SDS, 5% 2-mercaptoethanol, 0.0025% bromophenol blue). Immunoblotting was then performed with anti-Flag antibody. The samples were then sent to RIMUHC facility for mass spectrometry and proteomics analysis was done using the computational pipeline Scaffold.

## ACKNOWLEDGMENTS

We would like to thank Drs. Wendy Gilbert (Yale School of Medicine), Roy Parker (University of Boulder, Colorado), and Arlen Johnson (The University of Texas at Austin) for providing yeast strains, Kyla Edelmeier, Delina Efrem, Luca Lazzari, Ryan Huang, Suleima Jacob-Tomas, Sonya Madan, and Manassa Koripalli for technical help, and Dr. Joaquin Ortega (McGill University) for the localization of proteins in the structure of the 40S monosome and disomes (**Figure 6H**). Lokha R. Alagar Boopathy is funded by the FRQS scholarship program (315076) and the CRBS center at McGill University, Dr. Celia Alecki is funded by the FRQS postdoctoral fellowship (300232). This work is supported by NSERC grant RGPIN-2019-04767 and FRQ-NT NC-2999446 to Maria Vera.

## SUPPLEMENTARY INFORMATION

**Supplementary Figure 1.**
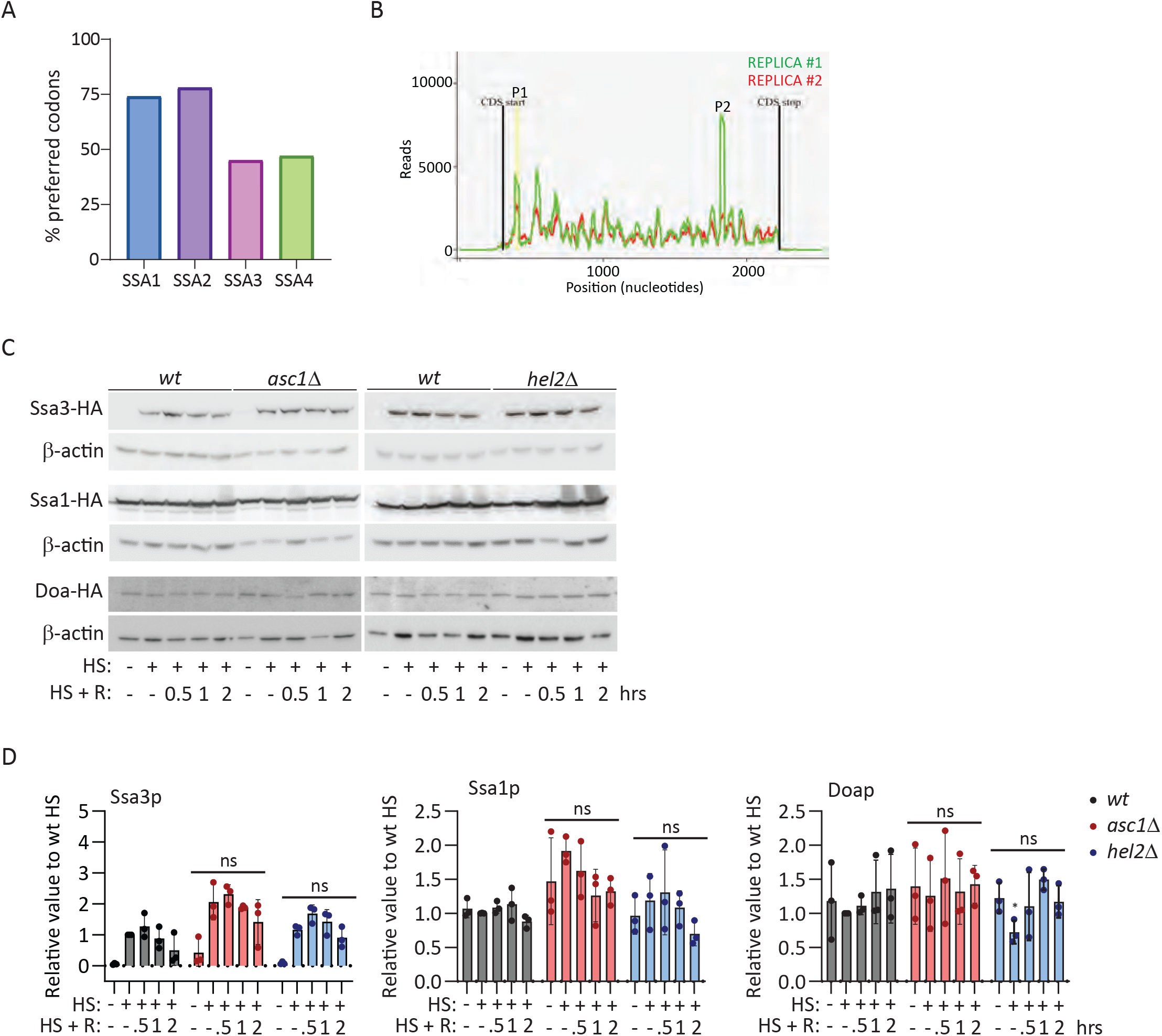
Asc1p and Hel2p do not regulate the translation of *SSA3* and *SSA1* mRNAs. **A.** Percentage of preferred codons in individual *SSA* mRNAs in *S.cerevisiae*: *SSA1* 74%, *SSA2* 78%, *SSA3* 45%, *SSA4* 47%. List of preferred codons in *S.cerevisiae* is procured from Bennetzen & Hall, 1982. Number indicates times a preferred codon is substitute to a low-frequent codon in *SSA4* mRNA **B.** Ribosome profiling analysis of *SSA4* mRNA in *Saccharomyces cerevisiae* upon 30 min of HS from Mühlhofer et al. 2019. Two independent replicates of ribosome density reads on the *SSA4* mRNA obtained from pausepred analysis (indicated in green and red) found 2 major ribosome pause sites P1 and P2 within the CDS to position 400 and 1831 in replica 1 and only P1 in replica 2. **C.** Immunoblots to detect the expression of Ssa3p (top), Ssa1p (middle) and Doap (bottom) tagged with 3 tandem HA epitopes and β-actin as the loading control in *wt, asc1Δ*, and *hel2Δ* strains under basal under basal, 30 min of HS at 42°C HS, and indicated recovery time points. **D.** Quantification of the expression of Ssa3p, Ssa1p and Doap. The HA band intensity in Ssa3p, Ssa1p, and Doap (right) was first normalized to the β-actin band intensity for each condition and then related to the normalized expression of HS *wt* yeast. The bars indicate the mean and SD of 3 independent experiments, each of them represented by a dot. Unpaired t-test (ns = non-significant differences, * = p<0.05).

**Supplementary Figure 2.**
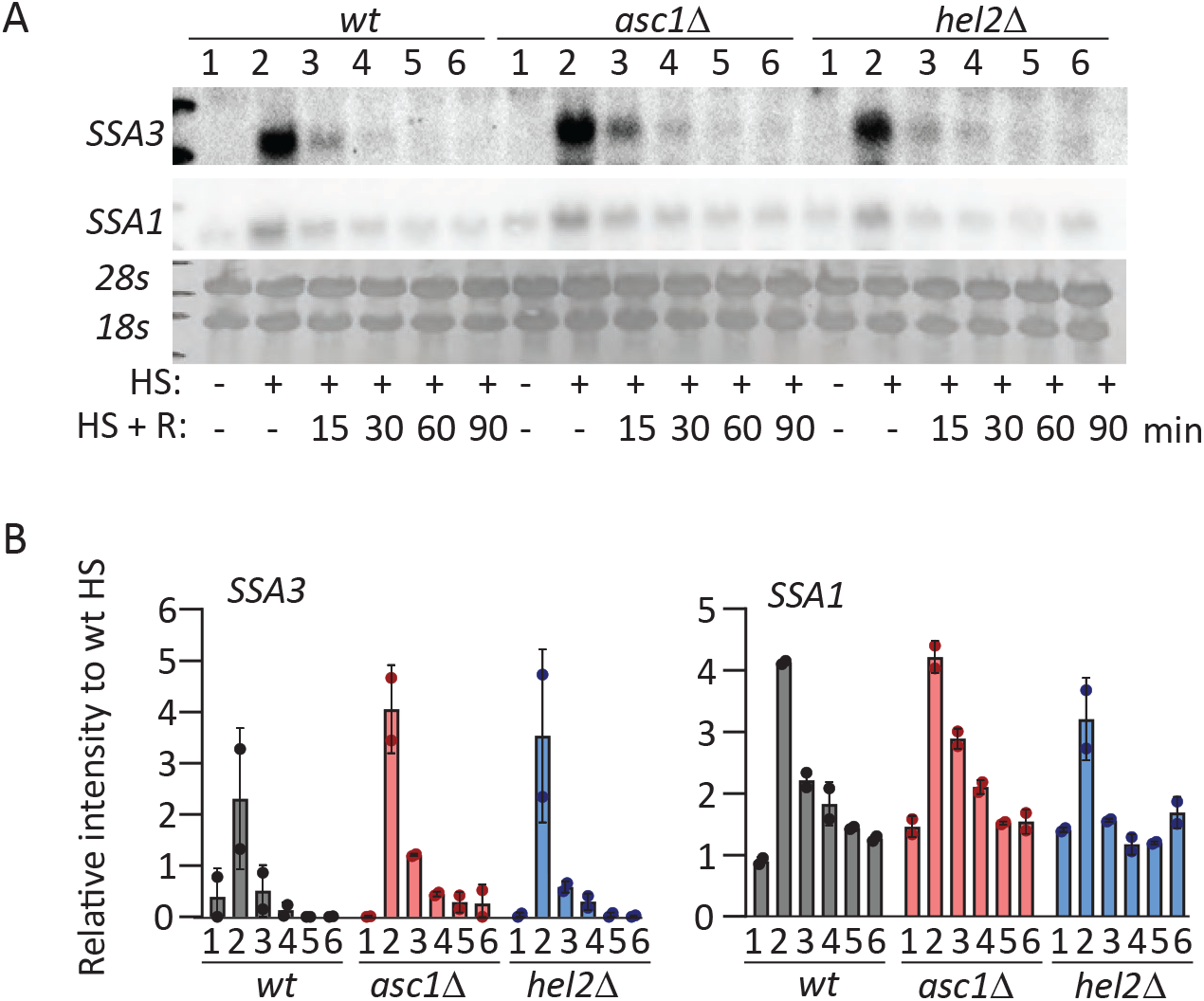
Asc1p and Hel2p do not stabilize SSA3 and SSA1 mRNAs during recovery. **A**. Northern blots to detect the expression of *SSA3* and *SSA1* mRNAs in *wt*, *asc1Δ*, and *hel2Δ* strains under basal, 30 min of HS at 42°C HS, and indicated recovery time points and corresponding methylene blue staining (bottom). **B**. Quantification of the expression of *SSA* mRNAs. The band intensity of *SSA3* and *SSA1* mRNAs was normalized to the RNA loading by the methylene blue staining. Bars indicate the average and SD of 2 independent experiments. No significant differences were found between *wt* and the *asc1Δ*, and *hel2Δ* strains.

**Supplementary Figure 3.**
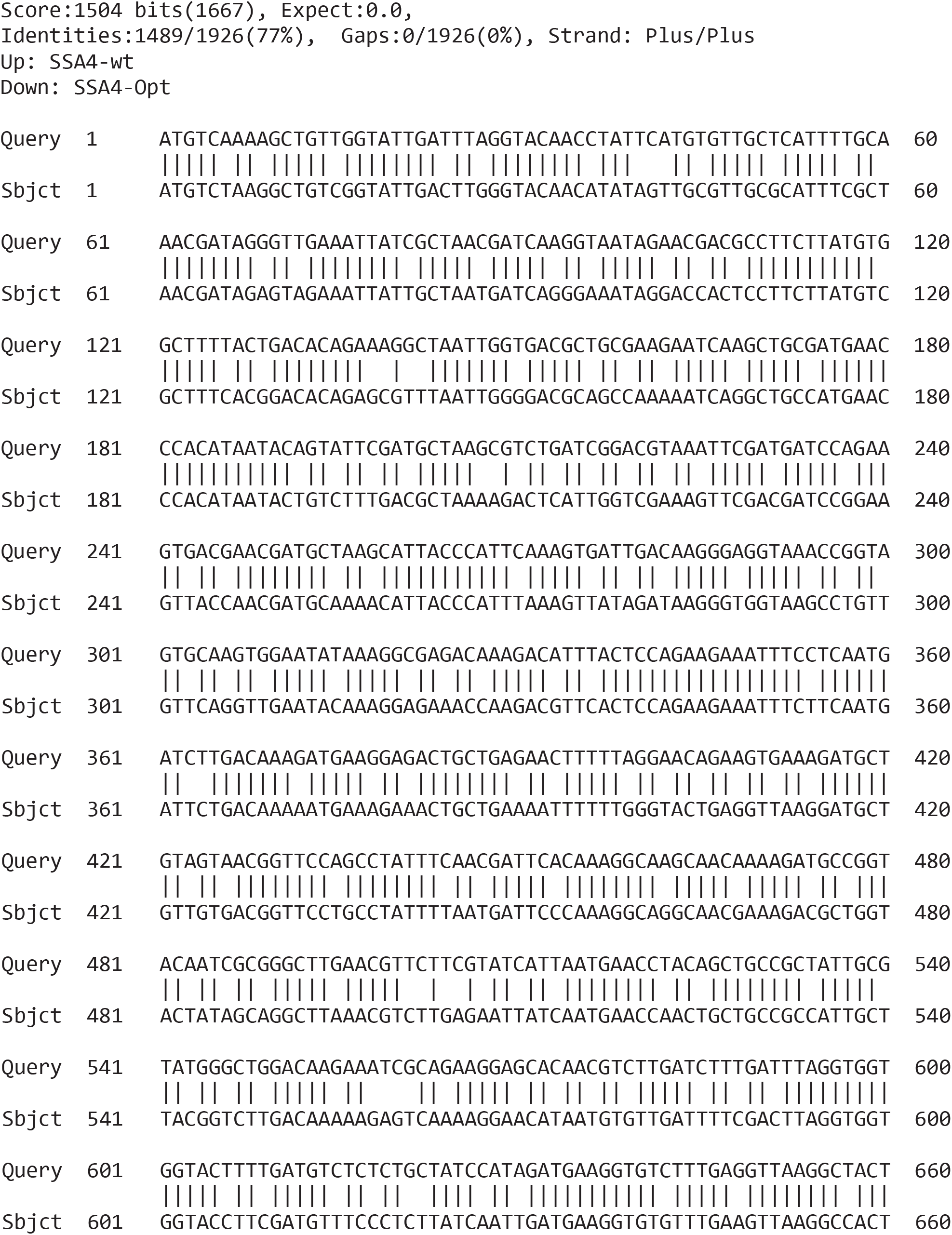
Codon optimization of SSA4 mRNA. Aligned of SSA4-wt (query) and SSA4 optimized (subject) coding sequences both containing 1926 nucleotides and sharing 77% identity (1487/1926).

**Supplementary Figure 4.**
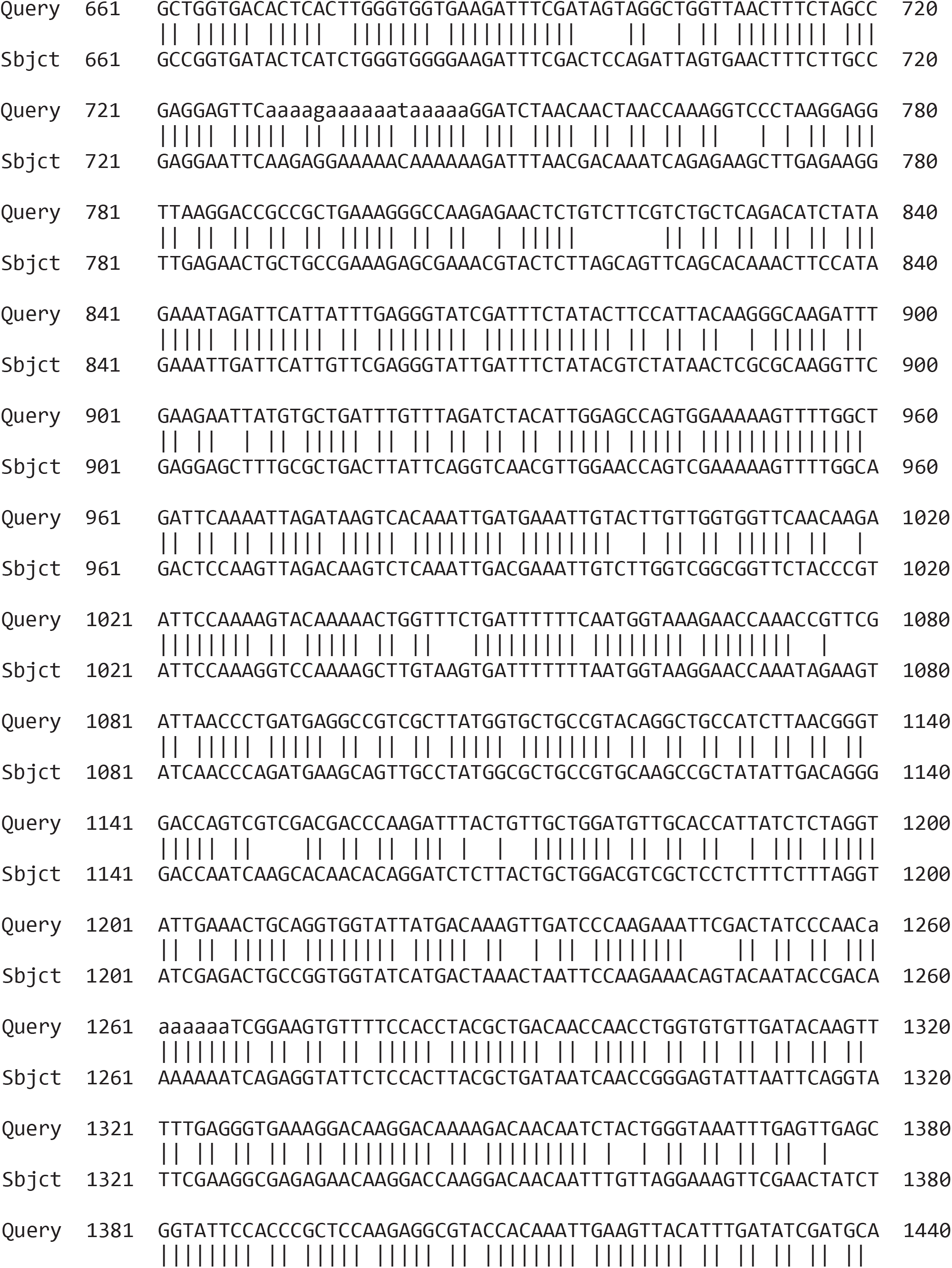
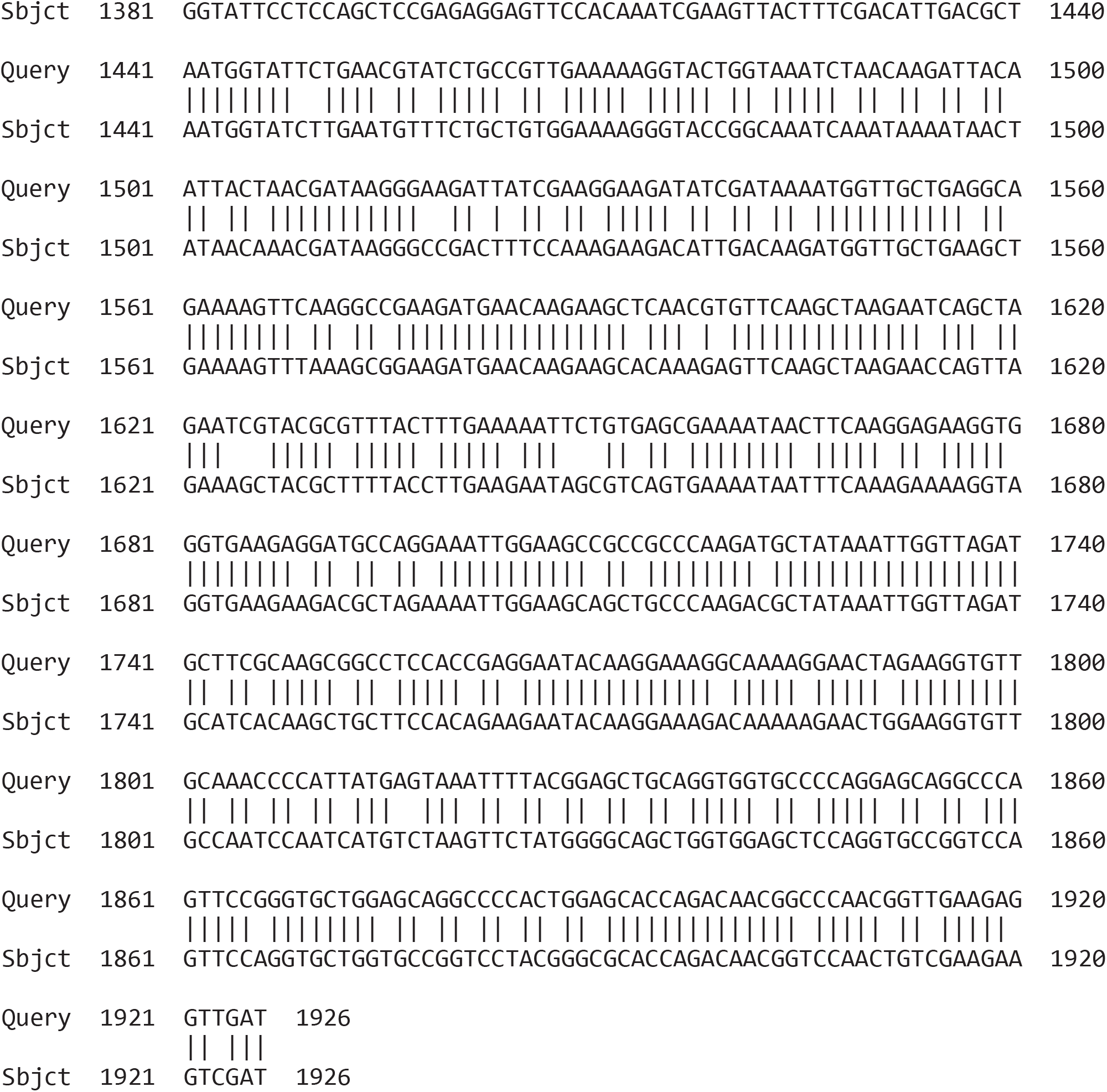
Asc1p regulates the Opt-SSA4 mRNA stability not SNR24 and no effect on Opt-SSA4 mRNA translation. **A.** RT-PCR to confirm the expression of *U24* (left), β*-actin* (middle) and Asc1 (right) in *wt*, *asc1Δ*, *asc1Δ* transform with a centromeric plasmid to express the CDS of Asc1p (*asc1Δ* + Asc1p), or *U24* from the intron of the *β-ACTIN* gene *(asc1Δ* + *U24*). **B.** Northern blots to detect the expression of *SSA4 Opt* (left) and *SSA2* (right) mRNAs in *Opt*, *Opt-ΔAsc1*, *Opt-ΔAsc1* with exogenous expression Asc1 (*Opt-asc1Δ*+ Asc1p), *Opt-asc1Δ* with exogenous expression of U24 *(Opt-asc1Δ* +*U24*) under basal, 30 min of HS at 42°C HS, and indicated recovery time points. (**C**) Quantification of the half-life of *SSA4 Opt* (left) and *SSA2* (right) mRNAs during recovery. The band intensity of *SSA4 Opt* and *SSA2* mRNA was normalized to the RNA loading by the methylene blue staining and then, each recovery time was related to the intensity of the HS (considered as 100% of induction) band for each strain to obtain the decay curve and calculate the half-life. **D.** Immunoblots to detect the expression of Ssa4p tagged with three tandem HA epitopes and Tubulin as the loading control *Opt*, *Opt-ΔAsc1* with exogenous expression Asc1 (*Opt-asc1Δ*+*Asc1p*), *Opt-asc1Δ*, *Opt-asc1Δ* with exogenous expression of U24 *(Opt-asc1Δ*+*U24*) yeast strains under basal, 30 min of HS at 42°C HS, and indicated recovery time points. **E.** Quantification of the expression of Ssa4p. The HA band intensity in ssa4 tagged strains was first normalized to the tubulin band intensity for each condition and then related to the normalized expression of *Opt-ΔAsc1* yeast under HS. The bars indicate the mean and standard deviation (SD) of three independent experiments each of them represented by a dot. Unpaired t-test (ns = non-significant).

**Supplementary Figure 5.**
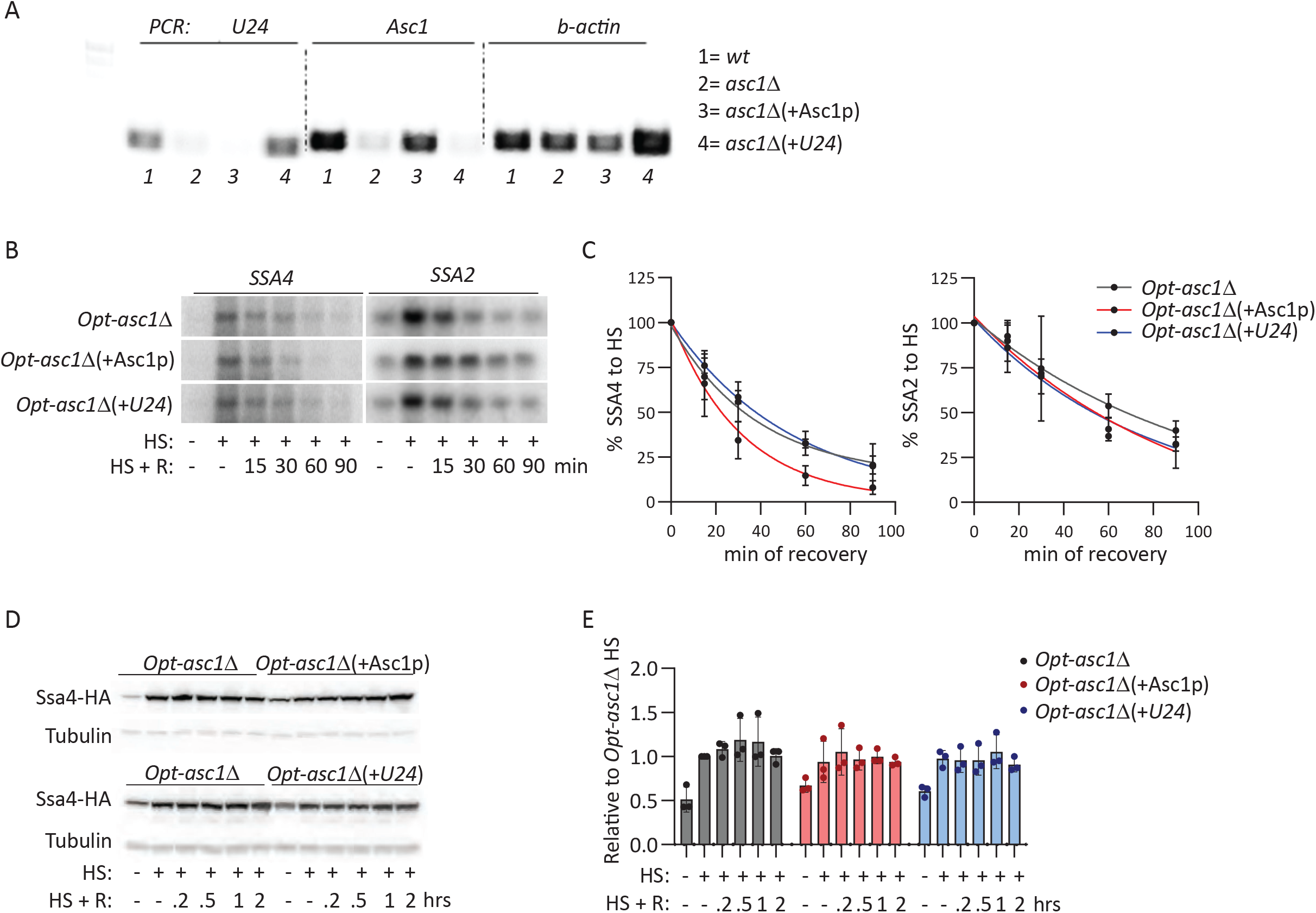
SSA4 mRNA degradation during recovery depends on Xrn1p and Dhh1p. **A.** Northern blots to detect the expression of *SSA4* (left) and *SSA2* mRNAs in *wt*, *not4Δ*, *ski7Δ, xrn1Δ*, and *xrn1Δ* strains under basal, 30 min of HS at 42°C HS, and indicated recovery time points. **B.** Quantification of the half-life of *SSA4* (left) and *SSA2* (right) mRNAs during recovery. The band intensity of *SSA4* and *SSA2* mRNA was normalized to the RNA loading by the methylene blue staining and then, each recovery time was related to the intensity of the HS (considered as 100% of induction) band for each strain to obtain the decay curve and calculate the half-life. **C.** Immunoblots to detect the expression of Ssa4p tagged with three tandem HA epitopes and Tubulin as the loading control in *wt*, *not4Δ*, *ski7Δ, xrn1Δ*, and *xrn1Δ* strains under basal, 30 min of HS at 42°C HS, and indicated recovery time points. **D.** Quantification of the expression of Ssa4p. The HA band intensity in Ssa4 tagged strains was first normalized to the tubulin band intensity for each condition and then related to the normalized expression of *wt* yeast under HS. The bars indicate the mean and standard deviation (SD) of three independent experiments, each of them represented by a dot. Unpaired t-test (ns = non-significant).

